# The blowfly *Chrysomya latifrons* inhabits fragmented rainforests, but lacks genetic diversity and population structure

**DOI:** 10.1101/2022.01.12.476129

**Authors:** Nathan J. Butterworth, James F. Wallman, Nikolas P. Johnston, Blake M. Dawson, Angela McGaughran

## Abstract

Climate change and deforestation are causing rainforests to become increasingly fragmented, placing them at heightened risk of biodiversity loss. Invertebrates constitute the greatest proportion of this biodiversity, yet we lack basic knowledge of their population structure and ecology. It is not currently feasible to assess the population structure of every invertebrate species, so there is a compelling need to identify ‘indicator species’ that are broadly indicative of habitat-level patterns and processes. Blowflies are an ideal candidate, because they are widespread, abundant, and can be easily collected within rainforests.

Here, we present the first study of the blowfly *Chrysomya latifrons*, which is endemic to the rainforests of New South Wales, Australia. We genotyped 188 flies from 15 isolated rainforests and found low overall genetic diversity and a complete lack of genetic structure between populations – suggesting the presence of a single large panmictic population along ~1,000 km of the Australian east coast. This highlights that: (1) *Ch. latifrons* inhabits every rainforest in NSW and undoubtedly plays an important role in these ecosystems, but low genetic diversity may cause it to struggle to adapt to a changing climate; (2) strongly dispersing insects have the capacity to migrate between isolated rainforests, likely carrying pollen, parasites, phoronts, and pathogens with them to form crucial trophic networks; and (3) there is an urgent need for similar studies on poorly dispersing rainforest insects, as these may be the most fragmented and at highest risk of local extinction.

## Introduction

Due to climate change and deforestation, rainforests are becoming increasingly fragmented islands of endemic biodiversity (Bowman 2000; Lens et al. 2002; Laurance et al. 2017) that are highly susceptible to biodiversity loss (Williams et al. 2003; Mariani et al. 2019; Nolan et al. 2020). To make informed conservation decisions for rainforests and their inhabitants, we must understand how rainforest habitats are fragmented, and how this affects the genetic patterns and ecological processes of their flora and fauna.

Species that inhabit fragmented rainforests can vary significantly in their extent of population genetic structure and diversity – spanning a continuum from well-connected to fragmented populations with high to low genetic diversity (Leung et al. 1994; Brown et al. 2004; Milá et al. 2009; Brito and Arias 2010; Woltmann et al. 2012; Sadanandan and Rheindt 2015). These species-specific genetic responses to fragmentation result from the unique physiologies, dispersal capacities, resource requirements, and habitat continua of species, as well as the unique landscape structure of the fragments they inhabit (i.e., the presence and number of corridors between adjacent fragments) (Leung et al. 1994; Callens et al. 2011; Woltmann et al. 2012). As such, certain populations and species will suffer greater genetic consequences from habitat fragmentation than others and understanding these differences will enable identification of which populations and species may be at greatest risk of further habitat fragmentation and climate change.

While attention has focused on many vertebrate animals experiencing these anthropogenic threats (Shapcott 2000; Shoo et al. 2005; Mac Nally et al. 2009; Moritz et al. 2009; Martínez-Ramos et al. 2016), invertebrates have been largely neglected (Ellwood and Foster 2004; Snaddon 2008; Wardhaugh et al. 2012; Taylor et al. 2018; New 2018). Of all animals, invertebrates are the largest contributors to the function and biodiversity of rainforest ecosystems and are those at the greatest risk of local extinction (Legge et al. 2021; Marsh et al. 2021). Despite this, we have a very limited understanding of the population structure of most rainforest invertebrates (Radespiel and Bruford 2014), including whether there is gene flow among geographically disconnected fragments, and which species and populations require conservation priority. There is an urgent need to remedy this, as studies around the world are beginning to identify significant changes in invertebrate populations, with habitat fragmentation and climate change as key drivers (Cardoso et al. 2020; Samways et al. 2020; Marsh et al. 2021).

In Australia, rainforests are naturally fragmented, and hold a substantial proportion of the continent’s total invertebrate biodiversity (Kitching et al. 1993; Yeates et al. 2003; Stork and Grimbacher 2006). Invertebrates play a crucial role in maintaining the function of these ecosystems (Ewers et al. 2015; Griffiths et al. 2017) – acting as primary decomposers, pollinators, parasites, and trophic resources. It is therefore important to understand how invertebrates are distributed throughout rainforests, whether there is gene flow between populations, and whether populations are highly adapted to their environments and therefore less likely to cope with sudden changes to climate or habitat structure (Kelly 2019; Razgour et al. 2019). However, given the overwhelming diversity of rainforest invertebrates, it is not currently feasible to comprehensively assess the genetic responses of every species to habitat fragmentation. As such, there is a need for ‘indicator species’ that inhabit fragmented rainforests, but are broadly distributed throughout them, can easily disperse between them, and can be reliably collected. Such species should show generally high levels of genetic connectivity between habitat fragments, and any genetically isolated populations should reflect highly isolated habitat fragments.

Blowflies (Diptera: Calliphoridae) are perfectly suited to this indicator approach because they are widespread throughout rainforests, extremely abundant, and easy to capture in the wild (Norris 1959; Badenhorst and Villet 2018; Butterworth et al. 2020). Blowflies are also highly vagile (Norris 1959) and many species exhibit wide dietary breadths – opportunistically feeding on a range of resources, including vertebrate and invertebrate carrion, dung, decaying plant matter, and pollen (Dear 1985; Brodie et al. 2015). As such, blowflies are likely some of the most capable invertebrate dispersers and should show high levels of connectivity between fragmented habitats.

Here, we use genotype-by-sequencing (GBS) through the DarTseq™ platform to obtain genetic information (single nucleotide polymorphisms; ‘SNPs’), targeting a species that is endemic to southeast Australia – *Chrysomya latifrons* Malloch 1927 (Figure 1). Ubiquitous in fragmented rainforests across a large (~1,000 km) geographic range and an able disperser, *Ch. latifrons* should show high levels of gene flow and diversity, and limited genetic differentiation between adjacent rainforests. We expect to observe only broad patterns of isolation by distance, so any rainforest fragments inhabited by *Ch. latifrons* that do not meet these patterns will suggest high levels of habitat isolation. This species thus has the potential to inform baseline patterns of rainforest connectivity and genetic diversity, and to provide an indication of which rainforest fragments may require greatest conservation priority under a changing climate.

**Figure 1.**
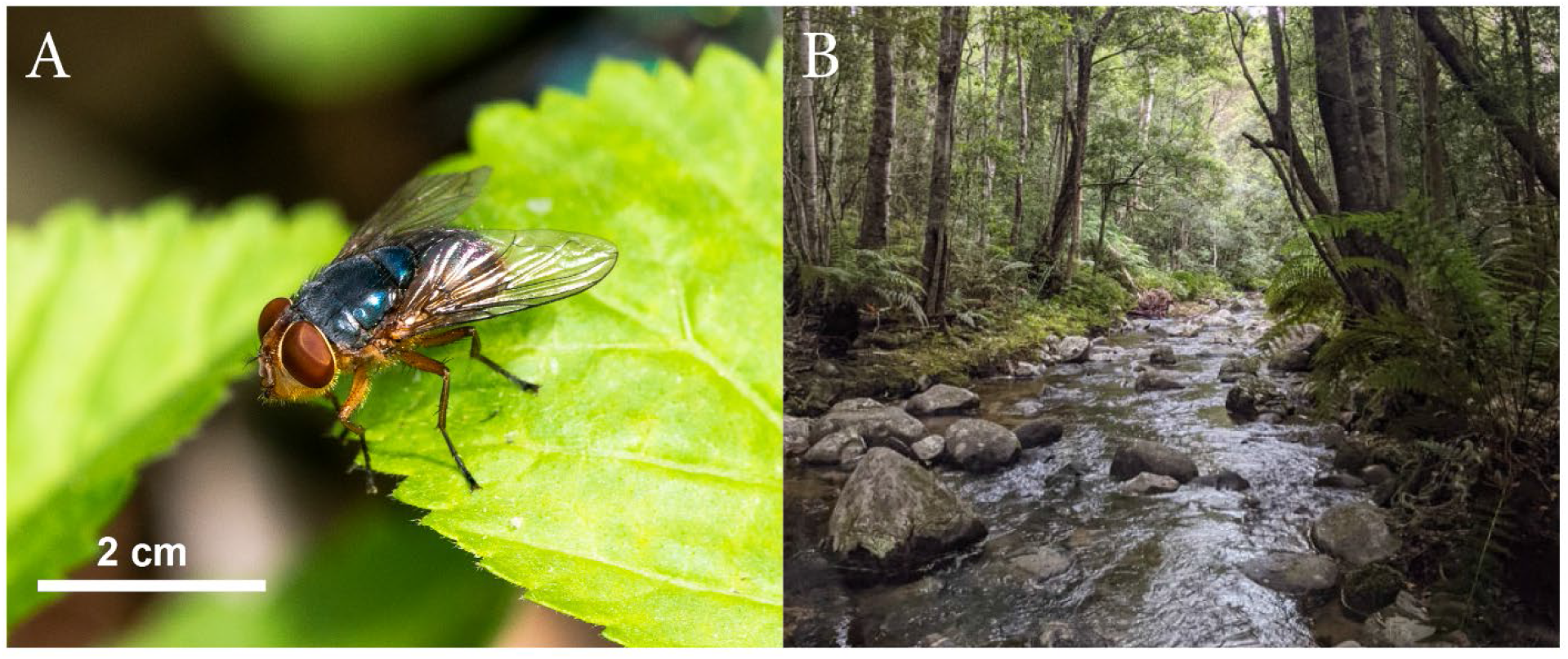
A: The blowfly *Chrysomya latifrons* which are endemic to the rainforests of southeast Australia; and B: the typical rainforest habitat where they can be found (pictured: Washpool National Park, NSW, Australia). Photographs taken by NB.

## Methods

### Specimen collecting

To create a bait that was attractive to *Ch. latifrons*, 500 g of raw kangaroo mince was left outdoors for 72 hours to attract *Calliphora* blowflies. Once *Calliphora* larvae had fed on the meat and reached the third instar, we tightly sealed the container for 24 hours. The subsequent anaerobic decomposition of *Calliphora* maggots (in combination with the decomposing kangaroo mince) encourages the growth of specific (but unknown) bacteria which emit volatiles that are highly attractive to all Australian *Chrysomya* species (except *Chrysomya flavifrons* Aldrich 1925). This bait presumably exploits the preference of *Chrysomya* species for cues associated with carrion that is in the mid-stage of decomposition, as most *Chrysomya* are secondary colonisers (Dawson et al. 2021).

Using this method, a total of 188 *Ch. latifrons* adults were caught with hand nets at locations between Washpool, NSW and Maxwells Rainforest, NSW (Table 1). Flies were euthanised with ethyl acetate vapour within eight hours of capture and placed into 2.5 mL plastic tubes containing 90% ethanol, then stored at −4°C in the laboratory for up to eight weeks. The head of each individual fly (3-18 from each population) was then dissected from the body, placed in a single well of a 96-well plate with 70% ethanol, and sent to Diversity Arrays Technology Pty Ltd (DarTseq™; Canberra, Australia; http://www.diversityarrays.com) for extraction and sequencing.

**Table 1.** Locality information, collection dates, and sample numbers from each of the sites where *Chrysomya latifrons* (Diptera: Calliphoridae) was collected. We also provide mean values of observed heterozygosity (Ho), expected heterozygosity (He), inbreeding coefficients (FIS) per population, and population specific FST (Weir and Goudet 2017).

| Site | Abbr. | Lat | Lon | Date | # | $H_o$ | $H_e$ | $F_{IS}$ | $F_{ST}$ |
| --- | --- | --- | --- | --- | --- | --- | --- | --- | --- |
| Barrington Tops National Park | BA | -32.1508 | 151.5248 | 27/01/21 | 16 | 0.180 | 0.212 | 0.129 | 0.000 |
| Bellangry | BE | -31.2892 | 152.5370 | 01/02/21 | 12 | 0.173 | 0.210 | 0.139 | 0.009 |
| Brown Mountain | BR | -36.5972 | 149.4440 | 18/02/21 | 10 | 0.171 | 0.206 | 0.126 | 0.032 |
| Foxground | FO | -34.6982 | 150.7920 | 26/11/20 | 12 | 0.173 | 0.211 | 0.144 | 0.007 |
| Goodenia Rainforest | GO | -36.8990 | 149.7154 | 20/02/21 | 16 | 0.175 | 0.213 | 0.138 | 0.000 |
| Grahams Trail | GR | -30.4245 | 152.8304 | 28/01/21 | 14 | 0.174 | 0.211 | 0.150 | 0.004 |
| Maxwells Rainforest | MA | -37.4145 | 149.8138 | 19/02/21 | 18 | 0.173 | 0.212 | 0.150 | 0.000 |
| Mt Cambewarra | CA | -34.8049 | 150.5722 | 07/12/20 | 12 | 0.173 | 0.210 | 0.137 | 0.009 |
| Mt Dromedary | DR | -36.2946 | 150.0337 | 17/02/21 | 17 | 0.183 | 0.212 | 0.114 | 0.000 |
| Mt Keira | KE | -34.4047 | 150.8713 | 24/11/21 | 12 | 0.177 | 0.210 | 0.124 | 0.008 |
| Ourimbah | OU | -33.3649 | 151.3939 | 26/01/21 | 12 | 0.177 | 0.209 | 0.118 | 0.013 |
| Pointer Gap | PO | -35.2605 | 150.3582 | 13/12/20 | 12 | 0.175 | 0.212 | 0.140 | 0.001 |
| Ulidarra National Park | UL | -30.2488 | 153.0855 | 31/01/21 | 3 | 0.174 | 0.210 | 0.086 | 0.039 |
| Washpool National Park | WA | -29.4700 | 152.3160 | 29/01/21 | 16 | 0.180 | 0.211 | 0.120 | 0.002 |
| Wyrabalong National Park | WY | -33.2938 | 151.5356 | 26/01/21 | 5 | 0.171 | 0.208 | 0.124 | 0.032 |

### DNA extraction and genotyping

Total genomic DNA was extracted from the heads of 188 flies at the DarTseq™ facility, following the DNA extraction protocol of Kilian et al. (2012). Genomic DNA was digested with the PstI-HpaII restriction enzyme pair, followed by PCR amplification, library construction, and Illumina Hiseq2500 sequencing (~1,200,000 reads per sample) as per the protocols of Kilian et al. (2012) and Georges et al. (2018). Following sequencing, reads were aligned to the genome of *Chrysomya rufifacies* Macquart 1843 (Andere et al. 2020), a close genetic relative of *Ch. latifrons* (Butterworth et al. 2020) with a sequenced genome. Genomic data were then processed using the DarTseq™ bioinformatic pipeline (Georges et al. 2018), which performed filtering and variant calling, and generated final genotypes.

### SNP filtering

Genotyping outputs were received from DArTseq™ in the DArT ‘2-row’ format – each allele is scored in a binary fashion where ‘1’ represents presence, ‘0’ represents absence, and ‘–’ represents a failure to score. The final dataset contained a total of 21,006 SNPs across 187 individual flies (Supplementary material 1). A single specimen was removed from the dataset (“L0303-Washpool_Male”) due to contamination. The data were then filtered in three steps with the ‘dartR’ package version 1.9.9.1 (Gruber et al. 2018) in R version 3.6.1 (R Core Team 2019). We first filtered the data by reproducibility (threshold: 0.98), then by call rate (threshold: 0.95), and finally by minor allele frequency (threshold: 0.02). This resulted in a filtered dataset of 187 individual genotypes, 2693 SNPs, and only 1.25% missing data, which was used for all downstream analyses.

### Genetic diversity

We used R for all analyses of genetic diversity. We first applied the ‘basic.stats’ function of the ‘hierfstat’ package version 0.5-10 (Goudet et al. 2021) to calculate average observed heterozygosity (Ho), expected heterozygosity (He), and inbreeding coefficients (F_IS_). We also used the ‘betas’ function from ‘hierfstat’ to calculate population-specific F_ST_ values (Weir and Goudet 2017).

### Population structure

Using R, we first assessed population structure by AMOVA using the function ‘poppr.amova’ with the ‘ade4’ implementation from the ‘poppr’ package version 2.9.3 (Kamvar et al. 2014). Then, to test whether populations were significantly different, we used a randomization test on the AMOVA output with 1,000 permutations (Excoffier et al. 1992) using the function ‘randtest’ from the package ‘ade4’ version 1.7-18 (Thioulouse et al. 2018). We then conducted pairwise comparisons of F_ST_ values between populations using the ‘gl.fst.pop’ function from the ‘dartR’ package with 10,000 bootstrap replicates.

Genetic distances between individuals were then examined using Nei’s distances, and a dendrogram with 1,000 bootstrap replicates was created with the ‘aboot’ function of the ‘poppr’ package, and the ‘ggtree’ function of the package ‘ggtree’ (Yu 2020). We then used the ‘glPca’ function from the ‘adegenet’ package version 2.1.5 (Jombart 2008) to determine whether genetic differences between individuals (as represented by principal components) were structured according to their populations.

To test for isolation by distance, we performed a Mantel test using the function ‘gl.ibd’ from the ‘dartR’ package in R. This compared linearised genetic distances between populations (calculated using ‘StAMPP’ version 1.6.3; Pembleton et al. 2013) against the Euclidean geographical distances (calculated using ‘vegan’ version 2.5-7; Oksanen et al. 2013).

To calculate individual blowfly admixture coefficients, the filtered SNP data were converted into the STRUCTURE format using the ‘gl2faststructure’ function from the ‘dartR’ package, then into the ‘.geno’ format using the ‘struct2geno’ function of the ‘LEA’ package version 3.1.4 (Frichot and Francois 2015). We then ran sparse non-negative matrix factorisation on these data with the ‘sNMF’ function from ‘LEA’. We analysed K values of 1 to 10, with 100 replications for each K value, and used the cross-entropy criterion to determine the value of K that best explained the results.

## Results

### Genetic diversity

Across all SNP loci, 75% of individuals were homozygous for the reference allele, 7% were homozygous for the alternate allele, and 18% were heterozygous – indicating relatively low genetic diversity across all populations. Low heterozygosity was also observed within each population, with population-specific Ho values all < 0.180, and all lower than the average expected heterozygosity (He) (Table 1). Further to this, more than 30% of loci had an average Ho of ≤ 0.1, and only 20% of loci had an average Ho > 0.3 (Supplementary material 2: Figure 1). Despite Ho being less than He in all populations, no signatures of inbreeding were detected, with low F_IS_ values in each population (always < 0.150).

### Population structure

Population specific F_ST_ values were low (ranging from 0.000 to 0.032), suggesting minimal genetic differentiation between populations. AMOVA was conducted to assess the partitioning of genetic variation within individuals, between individuals, and between populations (Table 2). Most of the variation was observed within individuals (83.5%) followed by between individuals (16.5%), and both were found to be significant factors contributing to the overall population structure (P < 0.01). However, genetic variation between populations was not detected (0.00%), consistent with the genetic diversity results and indicating that there is limited genetic differentiation between populations.

**Table 2.**
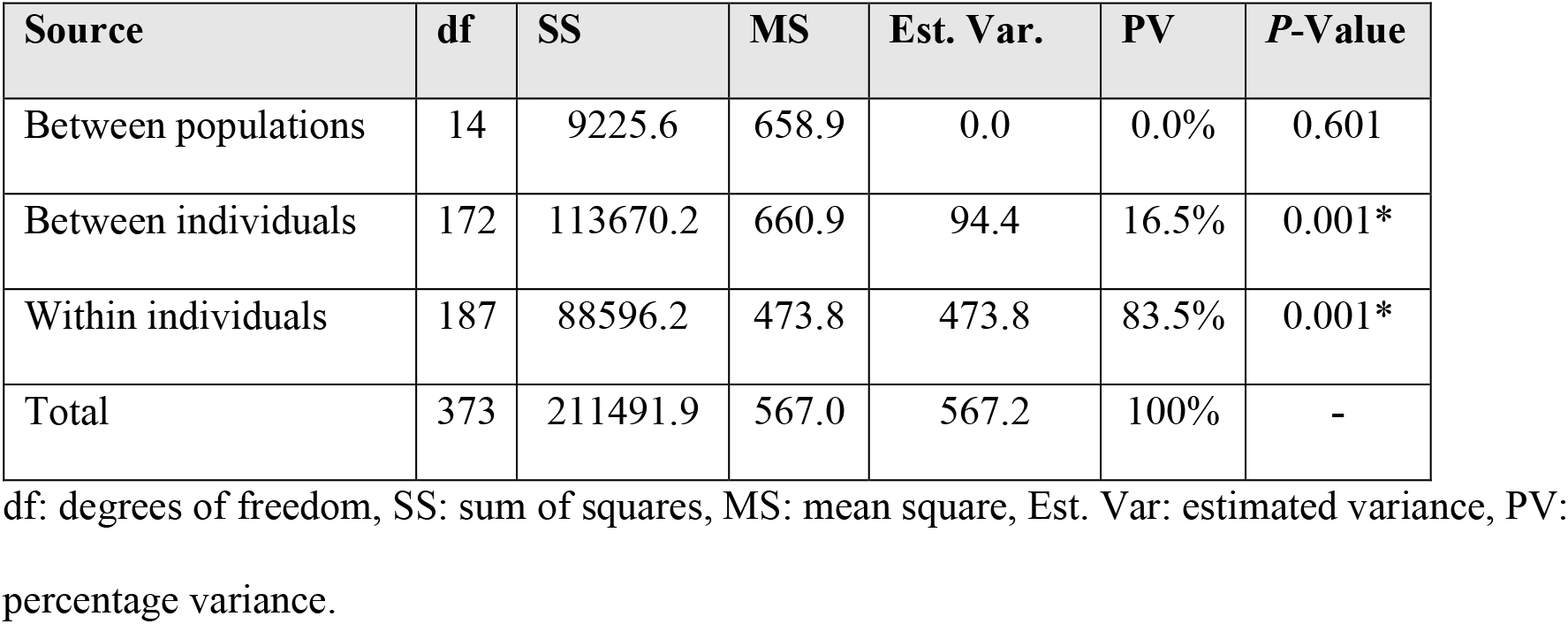
Analysis of molecular variance (AMOVA) to assess the extent of variation within and between 15 populations of *Chrysomya latifrons* (Diptera: Calliphoridae).

| Source | df | SS | MS | Est. Var. | PV | P-Value |
| --- | --- | --- | --- | --- | --- | --- |
| Between populations | 14 | 9225.6 | 658.9 | 0.0 | 0.0% | 0.601 |
| Between individuals | 172 | 113670.2 | 660.9 | 94.4 | 16.5% | 0.001* |
| Within individuals | 187 | 88596.2 | 473.8 | 473.8 | 83.5% | 0.001* |
| Total | 373 | 211491.9 | 567.0 | 567.2 | 100% | - |

df: degrees of freedom, SS: sum of squares, MS: mean square, Est. Var: estimated variance, PV: percentage variance.

Observed pairwise F_ST_ values were all < 0.009, and there were no significant differences detected between populations (Table 3).

**Table 3.** Pairwise FST values between 15 populations of *Chrysomya latifrons* (Diptera: Calliphoridae). Population abbreviations are provided in Table 1.

|  | KE | FO | CA | PO | BA | UL | BE | OU | WY | GR | WA | DR | BR | MA | GO |
| --- | --- | --- | --- | --- | --- | --- | --- | --- | --- | --- | --- | --- | --- | --- | --- |
| KE | - | - | - | - | - | - | - | - | - | - | - | - | - | - | - |
| FO | 0.000 | - | - | - | - | - | - | - | - | - | - | - | - | - | - |
| CA | 0.000 | 0.002 | - | - | - | - | - | - | - | - | - | - | - | - | - |
| PO | 0.000 | 0.000 | 0.000 | - | - | - | - | - | - | - | - | - | - | - | - |
| BA | 0.000 | 0.000 | 0.000 | 0.001 | - | - | - | - | - | - | - | - | - | - | - |
| UL | 0.000 | 0.000 | 0.003 | 0.000 | 0.000 | - | - | - | - | - | - | - | - | - | - |
| BE | 0.000 | 0.000 | 0.001 | 0.000 | 0.000 | 0.000 | - | - | - | - | - | - | - | - | - |
| OU | 0.000 | 0.000 | 0.001 | 0.000 | 0.000 | 0.000 | 0.000 | - | - | - | - | - | - | - | - |
| WY | 0.002 | 0.002 | 0.008 | 0.001 | 0.000 | 0.000 | 0.000 | 0.003 | - | - | - | - | - | - | - |
| GR | 0.000 | 0.000 | 0.000 | 0.000 | 0.000 | 0.000 | 0.000 | 0.000 | 0.003 | - | - | - | - | - | - |
| WA | 0.000 | 0.000 | 0.001 | 0.002 | 0.000 | 0.000 | 0.000 | 0.000 | 0.005 | 0.000 | - | - | - | - | - |
| DR | 0.000 | 0.000 | 0.000 | 0.000 | 0.000 | 0.004 | 0.000 | 0.000 | 0.003 | 0.000 | 0.001 | - | - | - | - |
| BR | 0.001 | 0.000 | 0.001 | 0.000 | 0.001 | 0.005 | 0.000 | 0.003 | 0.005 | 0.000 | 0.000 | 0.001 | - | - | - |
| MA | 0.001 | 0.000 | 0.001 | 0.000 | 0.000 | 0.000 | 0.001 | 0.000 | 0.002 | 0.000 | 0.001 | 0.000 | 0.002 | - | - |
| GO | 0.000 | 0.000 | 0.000 | 0.000 | 0.000 | 0.000 | 0.000 | 0.000 | 0.002 | 0.000 | 0.002 | 0.001 | 0.001 | 0.000 | - |

There was also no clear differentiation between individuals from different populations based on Nei’s genetic distances. In fact, individuals from populations separated by > 900 km often showed greater genetic similarity to each other than to individuals that were collected from the same geographic population (Figure 2). Consistent with these results, the mantel test of genetic vs geographic distance yielded no significant correlation (Mantel’s r = −0.053, *P* = 0.685) (Supplementary material 2: Figure 2). Likewise, in the principal component analysis (Figure 3, Supplementary material 2: Figure 3), the first two principal components explained only 0.97% and 0.95% of the total variation, and there was no clear separation of populations based on individual principal component scores (Figure 3).

**Figure 2.**
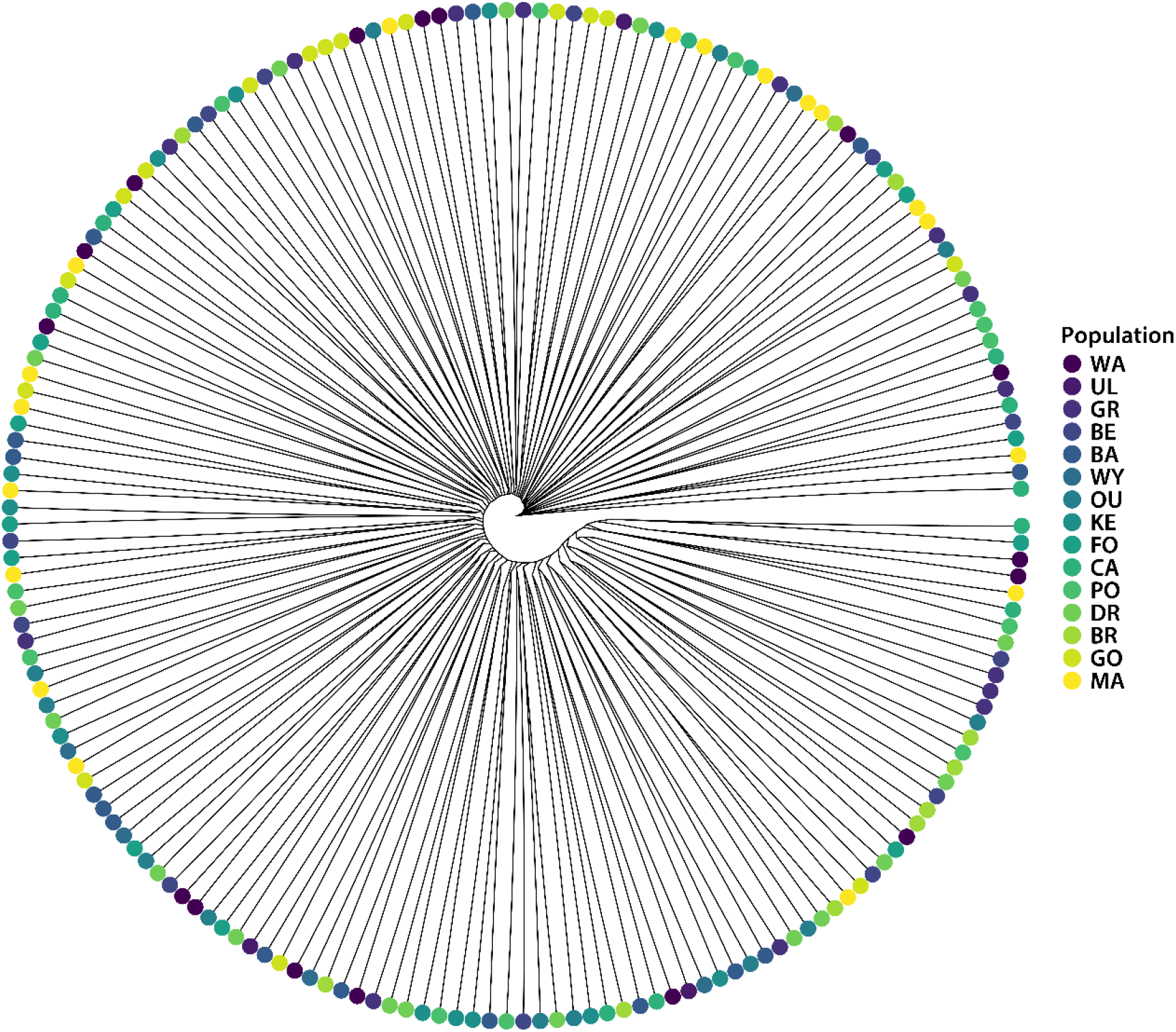
A dendrogram based on Nei’s genetic distances for individual *Chrysomya latifrons* (Diptera: Calliphoridae) from 15 rainforest populations. Populations are coloured according to the provided key in order from north (dark purple) to south (yellow). Population abbreviations are provided in Table 1.

**Figure 3.**
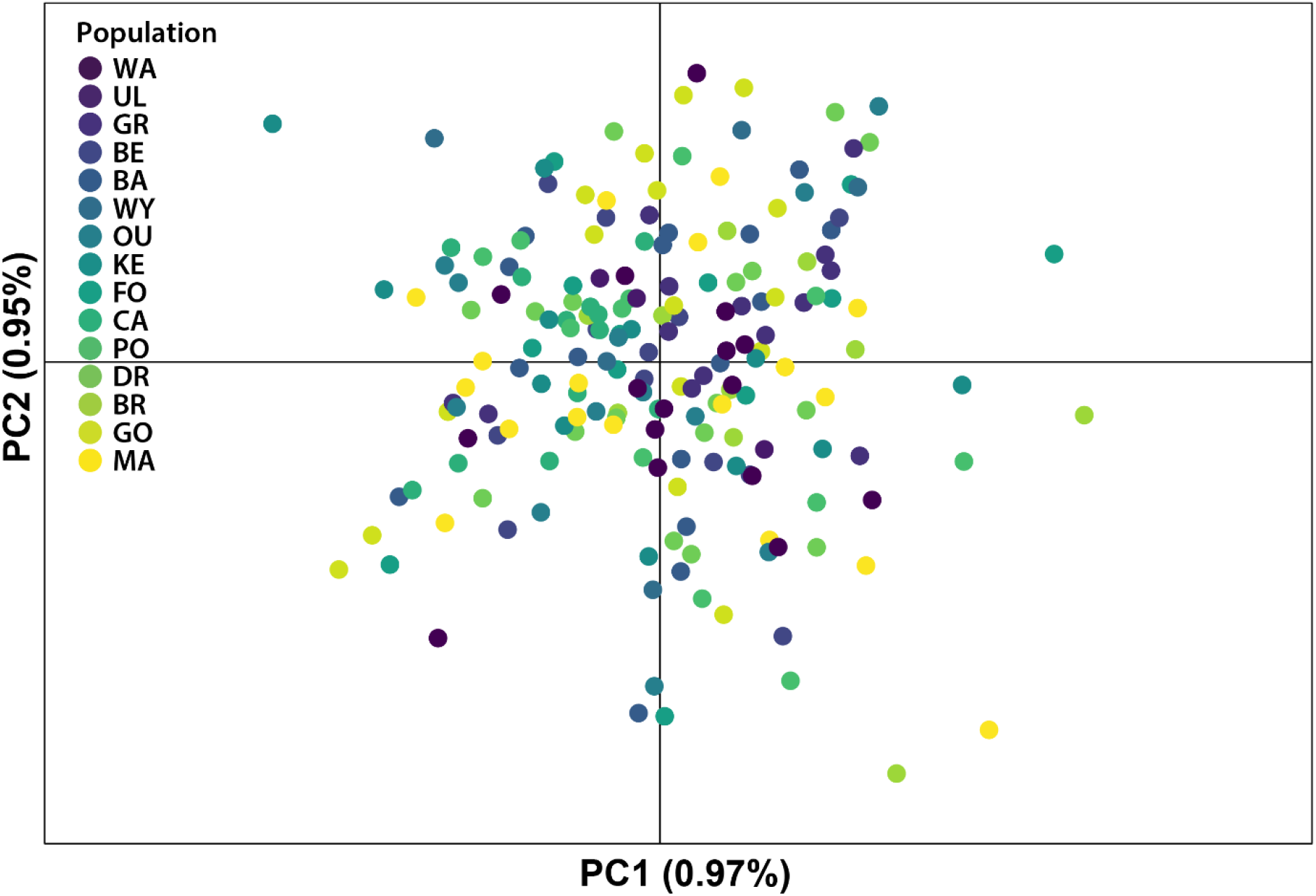
Principal component analysis of the filtered dataset of 2693 SNP loci from 15 populations of *Chrysomya latifrons* (Diptera: Calliphoridae). Populations are coloured according to the provided key in order from north (dark purple) to south (yellow) – following the same order as Figure 2. Population abbreviations are provided in Table 1.

The sNMF analysis identified the lowest cross-entropy for a K value of one (Supplementary material 2: Figure 4), which suggests that all populations share common ancestry. To investigate admixture in more detail, we explored the results for K-values from 1 to 10. For all tested K-values >1, we found that the admixture was evenly spread throughout geographic locations – supporting the notion that all populations share a common ancestry and that none were substantially genetically differentiated. To visualise these admixture proportions, we present the results of K=2 (Figure 4).

**Figure 4.**
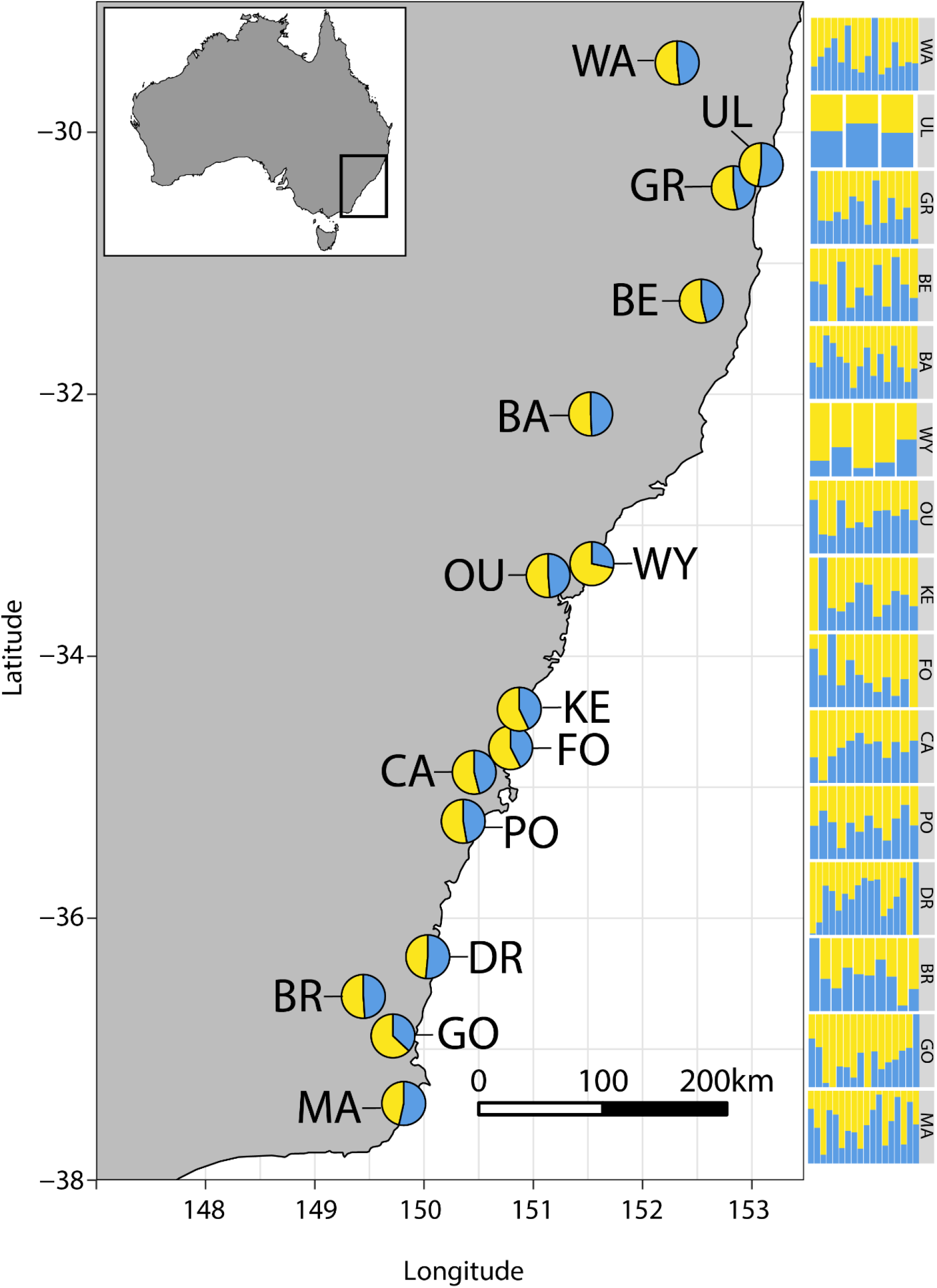
The mean admixture proportions of populations of *Chrysomya latifrons* (Diptera: Calliphoridae) that were sampled in the present study. The admixture proportions plotted on the map represent population averages. The bar plots presented on the right reflect individual admixture proportions, sorted by population, where each bar represents a single individual. Full population names are provided in Table 1. All analyses and plotting can be replicated by following the online tutorial provided by Tom Jenkins (https://github.com/Tom-Jenkins/admixture_pie_chart_map_tutorial).

## Discussion

We used the endemic Australian rainforest blowfly *Ch. latifrons* as an indicator species and genetic baseline for the analysis of connectivity among fragmented rainforests in Australia. Based on the ubiquity, widespread distribution, and high dispersal ability of *Ch. latifrons*, we expected to find high genetic diversity, limited genetic differences between adjacent rainforests, and only a broad pattern of isolation by distance. However, we found much lower than expected overall genetic diversity, and a complete lack of genetic structure between populations – indicative of a single large, genetically depauperate, panmictic population despite the ~1,000 km distribution span. Broadly, this highlights that strongly dispersing insects, such as blowflies, have the capacity to migrate between and connect isolated rainforest fragments – but that this does not preclude them from having low genetic diversity.

### Distribution

We found *Ch. latifrons* in high abundance from Washpool National Park in northern New South Wales to Maxwells Rainforest on the southern border with Victoria. This is the first comprehensive information on the distribution of *Ch. latifrons*, which inhabits a transect of > 1,000 km of eastern Australian rainforests. It is almost certain that populations also extend into the rainforests of northern Victoria, where there is suitable habitat available. Remarkably however, *Ch. latifrons* was completely absent near the Queensland border, where its closest genetic relative, *Ch. semimetallica*, occurs (Butterworth et al. 2020). These two species are rarely collected at the same location and likely compete with each other in a thin hybrid zone between Coffs Harbour in New South Wales and Brisbane in Queensland – providing an interesting model system for future work focused on understanding the eco-evolutionary dynamics of allopatry.

### Genetic diversity and population structure

Values of observed heterozygosity < 0.3 are generally considered to represent low genetic diversity (Robertson et al. 2018; Kleinhans and Willows-Munro 2019; Melo-Carrillo et al. 2020) and for all the populations sampled in the present study, the mean observed heterozygosity was < 0.18. In fact, more than 30% of loci had an observed heterozygosity of ≤ 0.1, and observed heterozygosity was consistently lower than expected heterozygosity. Together, this suggests a widespread lack of genetic diversity, despite these flies being in high abundance and spread over such a wide geographic range. In addition to this, population specific F_ST_ values indicate that population structure in *Ch. latifrons* is extremely low or absent, and this was reinforced by the consistent results of AMOVA, Nei’s genetic distances, principal component analysis, isolation-by-distance, and the sNMF analysis. Taken together, the diversity and differentiation results suggest that *Ch. latifrons* exists within east Australian rainforests as one large panmictic and genetically depauperate population. Considering that they exhibit such low genetic diversity and are endemic and restricted to New South Wales (Szpila and Wallman 2016), they are a good candidate for receiving conservation status.

The low genetic diversity and lack of population structure observed in *Ch. latifrons* highlights that even widely dispersing and abundant insects can be genetically depauperate and may be at risk of extinction under a changing climate. Comparably low levels of genetic variation and structure have been observed in many endangered vertebrate species, including the south African Cape vulture (Kleinhans and Willows-Munro 2019), the snail kite (Robertson et al. 2018), and the Mexican howler monkey (Melo-Carrillo et al. 2020). However, patterns of genetic connectivity and diversity are known to differ greatly between fly species – so not all species that inhabit fragmented habitats will be equally at risk of further fragmentation and climate change. For example, the tsetse fly *Glossina pallidipes* (which inhabits patchy and discontinuous habitat throughout Sub-Saharan Africa) shows clear population structure and high genetic diversity along a similar spatial scale to the present study (Bateta et al. 2020) and exhibits a high degree of variance in both desiccation resistance (Terblanche and Chown 2006) and thermal tolerance (Terblanche et al. 2008). Conversely, two species of Australian *Drosophila* (Diptera: Drosophilidae) (which inhabit fragmented rainforests) have been shown to have exceptionally low additive genetic variance and adaptability to desiccation resistance (Kellermann et al. 2006) and thermal stress (Saxon et al. 2017), suggesting they may have limited evolutionary potential to cope with climate change. Overall, it is unclear whether low genetic diversity and adaptive potential is a general pattern for flies living in fragmented habitats, but it is a question that requires rapid attention. As Australian rainforests are increasingly at risk of fire (Legge et al. 2021), it becomes more likely that their resident species will undergo further loss of (potentially already limited) genetic diversity.

### Biology

It is clear from our study that *Ch. latifrons* inhabits every major rainforest in south-eastern Australia in great abundances (for at least part of the year). This blowfly is therefore undoubtedly playing an important role in these ecosystems, but its biology is poorly understood. It is likely to be a carrion-breeding species, infesting vertebrate carrion in rainforests. This is supported by its attraction to carrion baits, oviposition on carrion, and the similarity of its larvae to other closely related carrion-breeding generalist species (Szpila and Wallman 2016). The widespread abundance and ability of *Ch. latifrons* to easily disperse long distances between habitats suggests they are either 1) generalist consumers of a wide range of animal carrion types, or 2) specialist consumers that utilise types of carrion that are abundant throughout rainforests.

The low population structure we observed in *Ch. latifrons* is unlikely to be a mere result of their strong dispersal ability, as there is ample evidence that flying insects with strong dispersal capacities often still show high levels of population structure (Bateta et al. 2020; Bluher et al. 2020). One possible explanation for the observed panmixia and low genetic diversity may relate to migration patterns. *Chrysomya* species tend to thrive in warmer temperatures (Byrd and Butler 1997; Sontigun et al. 2018), so it is plausible that a major source population is maintained in the warmer northern rainforests during winter, from which numerous individuals migrate towards southern rainforests over spring and summer.

This phenomenon is suggested to occur in Australian *Chrysomya rufifacies* (Norris 1959) and has also been reported for the blowfly *Cochliomyia hominivorax* in the Americas (Eddy and Bushland 1956). Blowflies have been shown to disperse as far as six km within 24 hours (Norris 1959), while some *Chrysomya* species can travel up to 65 km (Braack and Retief 1986), and individual stable flies (*Stomoxys calcitrans*) can migrate as far as 225 km (Hogsette and Ruff 1985). So, it is possible that long distance migration from a small winter source population explains the lack of population structure of *Ch. latifrons*. In fact, many Australian insects are known to migrate great distances and change distributions between seasons. For example, *Mythimna* and *Helicoverpa* moths (Lepidoptera) both make long south-easterly migrations in the warmer spring and summer months (Drake and Gatehouse 1995; Satterfield et al. 2020).

Strongly dispersing species with long-distance seasonal migration patterns generally tend to show low population structure over broad geographic ranges (Endersby et al. 2007; Pfeiler and Markow 2017; Wang et al. 2021), further supporting the possibility that large numbers of *Ch. latifrons* undertake south-easterly migrations during the spring and summer. There are numerous rainforest pockets and corridors that could facilitate these migrations, particularly along the mountainous Great Dividing Range, which extends from Queensland to Victoria.

However, our understanding of insect migration is still limited (Chapman et al. 2015; Pfeiler and Markow 2017; Satterfield et al. 2020) and a more comprehensive understanding is required, particularly as widespread distributions of insect populations in the summer months may mask the risk of local extinction of winter populations. This could occur in *Ch. latifrons*, particularly if a major source population is maintained in only a few northern NSW rainforests during the winter.

Physiological aspects may alternatively (or simultaneously) play a role in driving population structure in *Ch. latifrons*. Some *Chrysomya* species can cope with temperatures as low as 11°C (Cammack and Nelder 2010), and *Ch. latifrons* can occur around the greater Sydney region during the winter (Kavazos and Wallman 2012; Dawson et al. 2021). It may also be possible for individuals to diapause in southern regions throughout the winter (Saunders and Hayward 1998; Vinogradova and Reznik 2013), although there is no current evidence that *Ch. latifrons* has this capacity. Nevertheless, the temperature and climatic differences between northern and southern rainforests during winter may not be drastic enough to prevent *Ch. latifrons* from inhabiting southern rainforests throughout the entire year, meaning that migration may not be the main factor driving panmixia. Instead, panmixia may be the result of more continuous patterns of breeding between rainforests along eastern Australia. A lack of genetic diversity and population structure has also been shown along a north-south transect of Queensland fruit fly populations (Popa-Báez et al. 2020), supporting the idea that temperature variation along south-eastern Australia may not markedly constrain fly movement. There is still much to learn about how fly distributions and abundances change over the seasons in Australia; such knowledge will be instrumental to understanding the functioning of the continent’s ecosystems.

### Implications for rainforest connectivity

The movement of *Ch. latifrons* between isolated rainforests makes it a likely vector of dispersal for pollen, pathogens, parasites, and phoronts – forming an interconnected ecological network throughout southeast Australian rainforests. As such, we may also expect to see high degrees of genetic connectivity between the rainforest taxa that are associated with *Ch. latifrons*. Such correlated patterns of dispersal between fragmented habitats have been shown for rainforest trees and the African honeybees that pollinate them (Dick et al. 2003). Future work should therefore target the ecology of *Ch. latifrons*, particularly to understand their broad contributions to rainforest ecosystems and their adaptive potential in a changing climate.

Broadly, our results provide important insights into broad patterns of rainforest connectivity on the Australian continent – highlighting that rainforest invertebrates with strong dispersal capacities can move between, and connect, highly fragmented habitats. Such widespread dispersal between rainforests is, however, unlikely to be representative of all strong dispersers, depending largely on species-specific niche specialisations and trophic resources, which may constrain certain species to specific rainforest patches (e.g., Woltmann et al. 2012).

### Conclusions

The future capacity of species to disperse between rainforests is likely to be hampered by further climate change and deforestation, particularly with the increased likelihood of future wildfires (Legge et al. 2021). Within Australia, the long-term effects of the devastating 2019-2020 bushfires on habitat fragmentation and population dynamics are yet to be fully seen, and the results we present here will act as a crucial reference point for understanding these dynamics. There is an urgent need to study the genetic structure of more poorly dispersing rainforest insects that are endemic to fragmented rainforests. These are likely to be highly adapted to their specific habitats, and perhaps at highest risk of local extinction under a changing climate. Potential candidates are millipedes, worms, snails, ticks, spiders, and springtails – all of which are widely dispersed through rainforests (Mesibov 1998; Olson 1994) but are completely lacking information on population genetic structure. We strongly encourage researchers to identify such target species to begin further understanding rainforest population dynamics and conservation priorities.

## Supporting information

Supplementary metadata

## Acknowledgements

The scientific licence number associated with this project is SL101850. The authors thank the Paddy Pallin Foundation and the Royal Zoological Society of New South Wales for their financial support. We thank Finlay Davidson for his assistance with field work in southern NSW. NB also thanks Martin Butterworth, Roslyn Butterworth, and Kathryn Doty for their support.

## Data availability statement

Raw genotypes and metadata are available as supplementary material and will be uploaded to the Data Dryad repository upon acceptance in an academic journal.

## References

Andere, A. A., Pimsler, M. L., Tarone, A. M., & Picard, C. J. (2020). The genomes of a monogenic fly: views of primitive sex chromosomes. Scientific Reports, 10, 15728. https://doi.org/10.1038/s41598-020-72880-0

Badenhorst, R., & Villet, M. H. (2018). The uses of Chrysomya megacephala (Fabricius, 1794) (Diptera: Calliphoridae) in forensic entomology. Forensic Sciences Research, 3, 2–15. https://doi.org/10.1080/20961790.2018.1426136

Bateta, R., Saarman, N. P., Okeyo, W. A., Dion, K., Johnson, T., Mireji, P. O., Okoth, S., Malele, I., Murilla, G., Aksoy, S., & Caccone, A. (2020). Phylogeography and population structure of the tsetse fly Glossina pallidipes in Kenya and the Serengeti ecosystem. PLOS Neglected Tropical Diseases, 14, e0007855. https://doi.org/10.1371/journal.pntd.0007855

Bierregaard, R. O., Lovejoy, T. E., Kapos, V., Angelo Augusto dos, S., & Hutchings, R. W. (1992). The biological dynamics of tropical rainforest fragments. BioScience, 42, 859–866. https://doi.org/10.2307/1312085

Bluher, S. E., Miller, S. E. & Sheehan, M. J. (2020). Fine-scale population structure but limited genetic differentiation in a cooperatively breeding paper wasp. Genome Biology and Evolution, 12, 701–714. https://doi.org/10.1093/gbe/evaa070

Bowman, D. M. J. S. (2000). Australian rainforests: islands of green in a land of fire, Cambridge University Press, United Kingdom.

Braack, L. E., & Retief, P. F. (1986). Dispersal, density and habitat preference of the blow-flies Chrysomyia albiceps (Wd.) and Chrysomyia marginalis (Wd.) (Diptera: Calliphoridae). Onderstepoort Journal of Veterinary Research, 53, 13–18.

Brito, R. M. & Arias, M. C. (2010). Genetic structure of Partamona helleri (Apidae, Meliponini) from Neotropical Atlantic rainforest. Insectes Sociaux, 57, 413–419. https://doi.org/10.1007/s00040-010-0098-x

Brodie, B. S., Smith, M. A., Lawrence, J. & Gries, G. (2016). Effects of floral scent, color and pollen on foraging decisions and oocyte development of common green bottle flies. PLOS ONE, 10, e0145055. https://doi.org/10.1371/journal.pone.0145055

Brown, L. M., Ramey, R. R., Tamburini, B. & Gavin, T. A. (2004). Population structure and mitochondrial DNA variation in sedentary Neotropical birds isolated by forest fragmentation. Conservation Genetics, 5, 743–757. https://doi.org/10.1007/s10592-004-1865-x

Butterworth, N. J., Wallman, J. F., Drijfhout, F. P., Johnston, N. P., Keller, P. A. & Byrne, P. G. (2020). The evolution of sexually dimorphic cuticular hydrocarbons in blowflies (Diptera: Calliphoridae). Journal of Evolutionary Biology, 33, 1468–1486. https://doi.org/10.1111/jeb.13685

Byrd, J. H. & Butler, J. F. (1997). Effects of temperature on Chrysomya rufifacies (Diptera: Calliphoridae) development. Journal of Medical Entomology, 34, 353–358. https://doi.org/10.1093/jmedent/34.3.353

Callens, T. O. M., Galbusera, P., Matthysen, E., Durand, E. Y., Githiru, M., Huyghe, J. R. & Lens, L. U. C. (2011). Genetic signature of population fragmentation varies with mobility in seven bird species of a fragmented Kenyan cloud forest. Molecular Ecology, 20, 1829–1844. https://doi.org/10.1111/j.1365-294X.2011.05028.x

Cammack, J. A. & Nelder, M. P. (2010). Cool-weather activity of the forensically important hairy maggot blow fly Chrysomya rufifacies (Macquart) (Diptera: Calliphoridae) on carrion in upstate South Carolina, United States. Forensic Science International, 195, 139–142. https://doi.org/10.1016/j.forsciint.2009.12.007

Cardoso, P., Barton, P. S., Birkhofer, K., Chichorro, F., Deacon, C., Fartmann, T., Fukushima, C. S., Gaigher, R., Habel, J. C., Hallmann, C. A., Hill, M. J., Hochkirch, A., Kwak, M. L., Mammola, S., Ari Noriega, J., Orfinger, A. B., Pedraza, F., Pryke, J. S., Roque, F. O., Settele, J., Simaika, J. P., Stork, N. E., Suhling, F., Vorster, C. & Samways, M. J. (2020). Scientists’ warning to humanity on insect extinctions. Biological Conservation, 242, 108426. https://doi.org/10.1016/j.biocon.2020.108426

Chapman, J. W., Reynolds, D. R. & Wilson, K. (2015). Long-range seasonal migration in insects: mechanisms, evolutionary drivers and ecological consequences. Ecology Letters, 18, 287–302. https://doi.org/10.1111/ele.12407

Dawson, B. M., Barton, P. S. & Wallman, J. F. (2021). Field succession studies and casework can help to identify forensically useful Diptera. Journal of Forensic Sciences, 66, 2319–2328. https://doi.org/10.1111/1556-4029.14870

Dear, J. P. (1985). Calliphoridae (Insecta: Diptera). Fauna of New Zealand, 8.

Dick, C. W., Etchelecu, G. & Austerlitz, F. (2003). Pollen dispersal of tropical trees (Dinizia excelsa: Fabaceae) by native insects and African honeybees in pristine and fragmented Amazonian rainforest. Molecular Ecology, 12, 753–764. https://doi.org/10.1046/j.1365-294X.2003.01760.x

Drake, V. A. & Gatehouse, A. G. (1995). Insect migration: tracking resources through space and time. United Kingdom, Cambridge University Press.

Eddy, G. W. & Bushland, R. (1956). Screwworms that attack livestock. Yearbook of Agriculture, 1956, 172–175.

Ellwood, M. D. F. & Foster, W. A. (2004). Doubling the estimate of invertebrate biomass in a rainforest canopy. Nature, 429, 549–551. https://doi.org/10.1038/nature02560

Endersby, N. M., Hoffmann, A. A., McKechnie, S. W. & Weeks, A. R. (2007). Is there genetic structure in populations of Helicoverpa armigera from Australia? Entomologia Experimentalis et Applicata, 122, 253–263. https://doi.org/10.1111/j.1570-7458.2006.00515.x

Ewers, R. M., Boyle, M. J. W., Gleave, R. A., Plowman, N. S., Benedick, S., Bernard, H., Bishop, T. R., Bakhtiar, E. Y., Chey, V. K., Chung, A. Y. C., Davies, R. G., Edwards, D. P., Eggleton, P., Fayle, T. M., Hardwick, S. R., Homathevi, R., Kitching, R. L., Khoo, M. S., Luke, S. H., March, J. J., Nilus, R., Pfeifer, M., Rao, S. V., Sharp, A. C., Snaddon, J. L., Stork, N. E., Struebig, M. J., Wearn, O. R., Yusah, K. M. & Turner, E. C. (2015). Logging cuts the functional importance of invertebrates in tropical rainforest. Nature Communications, 6, 6836. https://doi.org/10.1038/ncomms7836

Excoffier, L., Smouse, P. E., & Quattro, J. M. (1992). Analysis of molecular variance inferred from metric distances among DNA haplotypes: application to human mitochondrial DNA restriction data. Genetics, 131, 479–491. https://doi.org/10.1093/genetics/131.2.479

Ferrar, P. (1987). A guide to the breeding habits and immature stages of Diptera Cyclorrhapha. Leiden, The Netherlands, Brill.

Frichot, E. & François, O. (2015). LEA: An R package for landscape and ecological association studies. Methods in Ecology and Evolution, 6, 925–929. https://doi.org/10.1111/2041-210X.12382

Gbedevi, K. M., Boukar, O., Ishikawa, H., Abe, A., Ongom, P. O., Unachukwu, N., Rabbi, I. & Fatokun, C. (2021). Genetic diversity and population structure of cowpea (Vigna unguiculata (L.) Walp.) germplasm collected from Togo based on DArT markers. Genes, 12, 1451. https://doi.org/10.3390/genes12091451

Georges, A., Gruber, B., Pauly, G. B., White, D., Adams, M., Young, M. J., Kilian, A., Zhang, X., Shaffer, H. B. & Unmack, P. J. (2018). Genomewide SNP markers breathe new life into phylogeography and species delimitation for the problematic short-necked turtles (Chelidae: Emydura) of eastern Australia. Molecular Ecology, 27, 5195–5213. https://doi.org/10.1111/mec.14925

Goudet, J., Jombart, T. & Goudet, M. J. (2015) Package ‘hierfstat’. R package version 0.04-22. http://github.com/jgx65/hierfstat.

Griffiths, H. M., Ashton, L. A., Walker, A. E., Hasan, F., Evans, T. A., Eggleton, P. & Parr, C. L. (2018). Ants are the major agents of resource removal from tropical rainforests. Journal of Animal Ecology, 87, 293–300. https:// https://doi.org/10.1111/1365-2656.12728

Gruber, B., Unmack P. J., Berry, O.F. & Georges, A. (2018). Dartr: An r package to facilitate analysis of SNP data generated from reduced representation genome sequencing. Molecular Ecology Resources, 18, 691–699. https://doi.org/10.1111/1755-0998.12745

Hardy, H. G. (1940). Notes on the Australian Muscoidea, V. Calliphoridae. Proceedings of the Royal Society of Queensland, 51, 133–146.

Hague, M. T. J. & Routman, E. J. (2016). Does population size affect genetic diversity? A test with sympatric lizard species. Heredity, 116, 92–98. https://doi.org/10.1038/hdy.2015.76

Hayward, S. A. & Saunders, D. S. (1998). Geographical and diapause-related cold tolerance in the blow fly, Calliphora vicina. Journal of Insect Physiology, 44, 541–551. https://doi.org/10.1016/s0022-1910(98)00049-3

Hogsette, J. A. & Ruff, J. P. (1985). Stable fly (Diptera: Muscidae) migration in northwest Florida. Environmental Entomology, 14, 170–175. https://doi.org/10.1093/ee/14.2.170

Jombart, T. (2008). adegenet: a R package for the multivariate analysis of genetic markers. Bioinformatics, 24, 1403–1405. https://doi.org/10.1093/bioinformatics/btn129.

Kamvar, Z. N., Tabima, J. F. & Grünwald, N. J. (2014). Poppr: an R package for genetic analysis of populations with clonal, partially clonal, and/or sexual reproduction. PeerJ, 2, e281. https://doi.org/10.7717/peerj.281

Kavazos, C. R. J. & Wallman, J. F. (2012). Community composition of carrion-breeding blowflies (Diptera: Calliphoridae) along an urban gradient in south-eastern Australia. Landscape and Urban Planning, 106, 183–190. https://doi.org/10.1016/j.landurbplan.2012.03.002

Kellermann, V. & van Heerwaarden, B. (2019). Terrestrial insects and climate change: adaptive responses in key traits. Physiological Entomology, 44, 99–115. https://doi.org/10.1111/phen.12282

Kellermann, V. M., Heerwaarden, B. V., Hoffmann, A. A. & Sgro, C. M. (2006). Very low additive genetic variance and evolutionary potential in multiple populations of two rainforest Drosophila species. Evolution, 60, 1104–1108. https://doi.org/10.1111/j.0014-3820.2006.tb01187.x.

Kelly, M. (2019). Adaptation to climate change through genetic accommodation and assimilation of plastic phenotypes. Philosophical Transactions of the Royal Society B: Biological Sciences, 374, 20180176. https://doi.org/10.1098/rstb.2018.0176

Ketema, S., Tesfaye, B., Keneni, G., Amsalu Fenta, B., Assefa, E., Greliche, N., Machuka, E. & Yao, N. (2020). DArTSeq SNP-based markers revealed high genetic diversity and structured population in Ethiopian cowpea (Vigna unguiculata (L.) Walp) germplasms. PLOS ONE, 15, e0239122. https://doi.org/10.1371/journal.pone.0239122

Kilian, A., Wenzl, P., Huttner, E., Carling, J., Xia, L., Blois, H., Caig, V., Heller-Uszynska, K., Jaccoud, D., Hopper, C., Aschenbrenner-Kilian, M., Evers, M., Peng, K., Cayla, C., Hok, P. & Uszynski, G. (2012). Diversity arrays technology: a generic genome profiling technology on open platforms. Methods in Molecular Biology, 888, 67–89. https://doi.org/10.1007/978-1-61779-870-2_5

Kitching, R. L., Bergelson, J. M., Lowman, M. D., McIntyre, S. & Carruthers, G. (1993). The biodiversity of arthropods from Australian rainforest canopies: General introduction, methods, sites and ordinal results. Australian Journal of Ecology, 18, 181–191. https://doi.org/10.1111/j.1442-9993.1993.tb00442.x

Kitching, R. L. & Voeten, R. (1977). The larvae of Chrysomya incisuralis (Macquart) and Ch. (Eucompsomyia) semimetallica (Malloch) (Diptera: Calliphoridae). Journal of the Australian Entomological Society, 16, 185–190.

Kleinhans, C. & Willows-Munro, S. (2019). Low genetic diversity and shallow population structure in the endangered vulture, Gyps coprotheres. Scientific Reports, 9, 5536. https://doi.org/10.1038/s41598-019-41755-4

Laurance, W. F., Camargo, J. L. C., Fearnside, P. M., Lovejoy, T. E., Williamson, G. B., Mesquita, R. C. G., Meyer, C. F. J., Bobrowiec, P. E. D. & Laurance, S. G. W. (2018). An Amazonian rainforest and its fragments as a laboratory of global change. Biological Reviews, 93, 223–247. https://doi.org/10.1111/brv.12343

Legge, S., Woinarski, J. C. Z., Scheele, B. C., Garnett, S. T., Lintermans, M., Nimmo, D. G., Whiterod, N. S., Southwell, D. M., Ehmke, G., Buchan, A., Gray, J., Metcalfe, D. J., Page, M., Rumpff, L., van Leeuwen, S., Williams, D., Ahyong, S. T., Chapple, D. G., Cowan, M., Hossain, M. A., Kennard, M., Macdonald, S., Moore, H., Marsh, J., McCormack, R. B., Michael, D., Mitchell, N., Newell, D., Raadik, T. A. & Tingley, R. (2021). Rapid assessment of the biodiversity impacts of the 2019–2020 Australian megafires to guide urgent management intervention and recovery and lessons for other regions. Diversity and Distributions. https://doi.org/10.1111/ddi.13428.

Lens, L., Van Dongen, S., Norris, K., Githiru, M. & Matthysen, E. (2002). Avian persistence in fragmented rainforest. Science, 298, 1236–1238. https://doi.org/10.1126/science.1075664

Leung, L. K. P, Dickman, C. R., & Moore, A. L. (1994). Genetic variation in fragmented populations of an Australian rainforest rodent, Melomys cervinipes. Pacific Conservation Biology, 1, 58–65. https://doi.org/10.1071/PC930058

Mac Nally, R., Bennett, A. F., Thomson, J. R., Radford, J. Q., Unmack, G., Horrocks, G. & Vesk, P. A. (2009). Collapse of an avifauna: climate change appears to exacerbate habitat loss and degradation. Diversity and Distributions, 15, 720–730. https://doi.org/10.1111/j.1472-4642.2009.00578.x

Mariani, M., Fletcher, M. S., Haberle, S., Chin, H., Zawadzki, A. & Jacobsen, G. (2019). Climate change reduces resilience to fire in subalpine rainforests. Global Change Biology, 25, 2030–2042. https://doi.org/10.1111/gcb.14609

Marsh, J., Bal, P., Fraser, H., Umbers, K., Greenville, A., Rumpff, L., & Woinarski, J. (2021). Assessment of the impacts of the 2019-20 wildfires of southern and eastern Australia on invertebrate species. NESP Threatened species recovery hub project 8.3.1. Final report, Brisbane.

Martínez-Ramos, M., Ortiz-Rodríguez, I. A., Piñero, D., Dirzo, R. & Sarukhán, J. (2016). Anthropogenic disturbances jeopardize biodiversity conservation within tropical rainforest reserves. Proceedings of the National Academy of Sciences, 113, 5323–5328. https://doi.org/10.1073/pnas.1602893113

Melo-Carrillo, A., Dunn, J. C. & Cortés-Ortiz, L. (2020). Low genetic diversity and limited genetic structure across the range of the critically endangered Mexican howler monkey (Alouatta palliata mexicana). American Journal of Primatology, 82, e23160. https://doi.org/10.1002/ajp.23160

Mesibov, R. (1998). Species-level comparison of litter invertebrates at two rainforest sites in Tasmania. Tasforests, 10, 141–153.

Milá, B., Wayne, R. K., Fitze, P. & Smith, T. B. (2009). Divergence with gene flow and fine-scale phylogeographical structure in the wedge-billed woodcreeper, Glyphorynchus spirurus, a Neotropical rainforest bird. Molecular Ecology, 18, 2979–2995. https://doi.org/10.1111/j.1365-294X.2009.04251.x

Monica, K., Mark, D., John, C., Ninon, M. & Katarina, M. M. (2021). Genome-wide SNPs detect fine-scale genetic structure in threatened populations of squirrel glider Petaurus norfolcensis. Research Square, preprint. https://doi.org/10.21203/rs.3.rs-717093/v1.

Moritz, C., Hoskin, C., Graham, C., Hugall, A., & Moussalli, A. (2005). Historical biogeography, diversity and conservation of Australia’s tropical rainforest herpetofauna. In A. Purvis, J. Gittleman, & T. Brooks (Eds.), Phylogeny and Conservation (Conservation Biology, pp. 243–264). Cambridge: Cambridge University Press. https://doi.org/10.1017/CBO9780511614927.011

New, T. R. (2018). Forests and insect conservation in Australia. Cham, Switzerland. Springer international publishing.

Nolan, R. H., Boer, M. M., Collins, L., Resco de Dios, V., Clarke, H., Jenkins, M., Kenny, B. & Bradstock, R. A. (2020). Causes and consequences of eastern Australia’s 2019–20 season of mega-fires. Global Change Biology, 26, 1039–1041. https://doi.org/10.1111/gcb.14987

Norris, K. R. (1959). The ecology of sheep blowflies in Australia. In A. Keast, Crocker R. L. & Christian, C. S. (Eds). Biogeography and Ecology in Australia. Dordrecht, Springer Netherlands: 514–544.

Oksanen, J., Blanchet, F. G., Kindt, R., Legendre, P., Minchin, P. R., O’hara, R. B., Simpson, G. L., Solymos, P., Stevens, M. H., & Wagner, H. (2013). Package ‘vegan’. Community ecology package, version 2.5-7 https://CRAN.R-project.org/package=vegan

Olson, D. M. (1994). The distribution of leaf litter invertebrates along a Neotropical altitudinal gradient. Journal of Tropical Ecology, 10, 129–150. https://doi.org/10.1017/S0266467400007793

Pembleton, L., Cogan, N., & Forster, J. (2013). StAMPP: an R package for calculation of genetic differentiation and structure of mixed-ploidy level populations. Molecular Ecology Resources, 13, 946–952. https://doi.org/10.1111/1755-0998.12129

Pfeiler, E. & Markow, T. A. (2017). Population connectivity and genetic diversity in long-distance migrating insects: divergent patterns in representative butterflies and dragonflies. Biological Journal of the Linnean Society, 122, 479–486. https://doi.org/10.1093/biolinnean/blx074

Popa-Báez, Á. D., Catullo, R., Lee, S. F., Yeap, H. L., Mourant, R. G., Frommer, M., Sved, J. A., Cameron, E. C., Edwards, O. R., Taylor, P. W. & Oakeshott, J. G. (2020). Genome-wide patterns of differentiation over space and time in the Queensland fruit fly. Scientific Reports, 10, 10788. https://doi.org/10.1038/s41598-020-67397-5

Malloch, J. R. (1927). Notes on Australian Diptera. No. XI. Proceedings of the Linnean Society of New South Wales, 52, 299–335.

Radespiel, U. & Bruford, M. W. (2014). Fragmentation genetics of rainforest animals: insights from recent studies. Conservation Genetics, 15, 245–260. https://doi.org/10.1007/s10592-013-0550-3

Razgour, O., Forester, B., Taggart, J. B., Bekaert, M., Juste, J., Ibáñez, C., Puechmaille, S. J., Novella-Fernandez, R., Alberdi, A. & Manel, S. (2019). Considering adaptive genetic variation in climate change vulnerability assessment reduces species range loss projections. Proceedings of the National Academy of Sciences, 116, 10418. https://doi.org/10.1073/pnas.1820663116

Robertson, E. P., Fletcher, R. J. & Austin, J. D. (2018). Microsatellite polymorphism in the endangered snail kite reveals a panmictic, low diversity population. Conservation Genetics, 19, 337–348. https://doi.org/10.1007/s10592-017-1003-1

Sadanandan, K. R. & Rheindt, F. E. (2015). Genetic diversity of a tropical rainforest understory bird in an urban fragmented landscape. The Condor, 117, 447–459. https://doi.org/10.1650/CONDOR-14-199.1

Samways, M. J., Barton, P. S., Birkhofer, K., Chichorro, F., Deacon, C., Fartmann, T., Fukushima, C. S., Gaigher, R., Habel, J. C., Hallmann, C. A., Hill, M. J., Hochkirch, A., Kaila, L., Kwak, M. L., Maes, D., Mammola, S., Noriega, J. A., Orfinger, A. B., Pedraza, F., Pryke, J. S., Roque, F. O., Settele, J., Simaika, J. P., Stork, N. E., Suhling, F., Vorster, C. & Cardoso, P. (2020). Solutions for humanity on how to conserve insects. Biological Conservation, 242, 108427. https://doi.org/10.1016/j.biocon.2020.108427

Satterfield, D. A., Sillett, T. S., Chapman, J. W., Altizer, S. & Marra, P. P. (2020). Seasonal insect migrations: massive, influential, and overlooked. Frontiers in Ecology and the Environment, 18, 335–344. https://doi.org/10.1002/fee.2217

Shapcott, A. (2000). Conservation and genetics in the fragmented monsoon rainforest in the Northern Territory, Australia: a case study of three frugivore-dispersed species. Australian Journal of Botany, 48, 397–407. https://doi.org/10.1071/BT98081

Shoo, L. P., Williams, S. E. & Hero, J.-M. (2005). Climate warming and the rainforest birds of the Australian Wet Tropics: Using abundance data as a sensitive predictor of change in total population size. Biological Conservation, 125, 335–343. https://doi.org/10.1016/j.biocon.2005.04.003

Smith Thomas, B., Wayne Robert, K., Girman Derek, J. & Bruford Michael, W. (1997). A role for ecotones in generating rainforest biodiversity. Science, 276, 1855–1857. https://doi.org/10.1126/science.276.5320.1855

Snaddon, J. L., Turner, E. C. & Foster, W. A. (2008). Children’s perceptions of rainforest biodiversity: which animals have the lion’s share of environmental awareness? PLOS ONE, 3, e2579. https://doi.org/10.1371/journal.pone.0002579

Sontigun, N., Sukontason, K. L., Klong-klaew, T., Sanit, S., Samerjai, C., Somboon, P., Thanapornpoonpong, S. N, Amendt, J. & Sukontason, K. (2018). Bionomics of the oriental latrine fly Chrysomya megacephala (Fabricius) (Diptera: Calliphoridae): temporal fluctuation and reproductive potential. Parasites & Vectors, 11, 415. https://doi.org/10.1186/s13071-018-2986-2

Stork, N. E. & Grimbacher, P. S. (2006). Beetle assemblages from an Australian tropical rainforest show that the canopy and the ground strata contribute equally to biodiversity. Proceedings of the Royal Society B: Biological Sciences, 273, 1969–1975. https://doi.org/10.1098/rspb.2006.3521

Szpila, K. & Wallman, J. F. (2016). Morphology and identification of first instar larvae of Australian blowflies of the genus Chrysomya of forensic importance. Acta Tropica, 162, 146–154. https://doi.org/10.1016/j.actatropica.2016.06.006

Taylor, G. S., Braby, M. F., Moir, M. L., Harvey, M. S., Sands, D. P. A., New, T. R., Kitching, R. L., McQuillan, P. B., Hogendoorn, K., Glatz, R. V., Andren, M., Cook, J. M., Henry, S. C., Valenzuela, I. & Weinstein, P. (2018). Strategic national approach for improving the conservation management of insects and allied invertebrates in Australia. Austral Entomology, 57, 124–149. https://doi.org/10.1111/aen.12343

Terblanche, J. S. & Chown, S. L. (2006). The relative contributions of developmental plasticity and adult acclimation to physiological variation in the tsetse fly, Glossina pallidipes (Diptera, Glossinidae). Journal of Experimental Biology, 209, 1064–1073. https://doi.org/10.1242/jeb.02129

Terblanche, J. S., Clusella-Trullas, S., Deere, J. A. & Chown, S. L. (2008). Thermal tolerance in a south-east African population of the tsetse fly Glossina pallidipes (Diptera, Glossinidae): Implications for forecasting climate change impacts. Journal of Insect Physiology, 54, 114–127. https://doi.org/10.1016/j.jinsphys.2007.08.007

Thioulouse, J., Dray S., Dufour A., Siberchicot A., Jombart, T., & Pavoine, S. (2018). Multivariate analysis of ecological data with ade4. Springer, New York, NY. https://doi.org/10.1007/978-1-4939-8850-1

Vinogradova, E. B. & Reznik, S. Y. (2013). Induction of larval diapause in the blowfly, Calliphora vicina R.-D. (Diptera, Calliphoridae) under field and laboratory conditions. Entomological Review, 93, 935–941. https://doi.org/10.1134/S0013873813080010

Wang, X., Yang, X., Zhou, L., Wyckhuys, K. A. G., Jiang, S., Van Liem, N., Vi, L. X., Ali, A. & Wu, K. (2021). Population genetics unveils large-scale migration dynamics and population turnover of Spodoptera exigua. Pest Management Science, 78, 612–625. https://doi.org/10.1002/ps.6670

Wardhaugh, C. W., Stork, N. E., Edwards, W. & Grimbacher, P. S. (2012). The overlooked biodiversity of flower-visiting invertebrates. PLOS ONE, 7, e45796. https://doi.org/10.1371/journal.pone.0045796

Weir, B. S. & Goudet, J. (2017). A unified characterization of population structure and relatedness. Genetics, 206, 2085–2103. https://doi.org/10.1534/genetics.116.198424

Williams, S. E., Bolitho, E. E. & Fox, S. (2003). Climate change in Australian tropical rainforests: an impending environmental catastrophe. Proceedings of the Royal Society of London. Series B: Biological Sciences, 270, 1887–1892. https://doi.org/10.1098/rspb.2003.2464

Woltmann, S., Kreiser, B. R. & Sherry, T. W. (2012). Fine-scale genetic population structure of an understory rainforest bird in Costa Rica. Conservation Genetics, 13, 925–935. https://doi.org/10.1007/s10592-012-0341-2

Yeates, D., Harvey, M. & Austin, A. (2003). New estimates for terrestrial arthropod species-richness in Australia. Records of the South Australian Museum Monograph Series, 7, 231–241.

Yu, G. (2020). Using ggtree to visualize data on tree-like structures. Current Protocols in Bioinformatics, 69, e96. https://doi.org/10.1002/cpbi.96

