## Supplementary Material 2 for "The blowfly *Chrysomya latifrons* inhabits fragmented rainforests, but lacks genetic diversity and population structure"

### Blowflies as ecological indicators: *Chrysomya latifrons* inhabits fragmented rainforests but shows no population structure

Nathan J. Butterworth[1*], James F. Wallman[1], Nikolas P. Johnston[2], Blake M. Dawson[3], Angela McGaughran[4]

[1]Faculty of Science, University of Technology Sydney, Ultimo NSW 2007, Australia

[2]Department of Ecology and Biogeography, Faculty of Biological and Veterinary Sciences, Nicolaus Copernicus University in Toruń, 87-100 Toruń, Poland

[3]Centre for Sustainable Ecosystem Solutions, School of Earth, Atmospheric and Life Sciences, University of Wollongong, Wollongong, NSW, 2522, Australia

[4]Te Aka Mātuatua - School of Science, University of Waikato, Private Bag 3105, Hamilton 3240, New Zealand

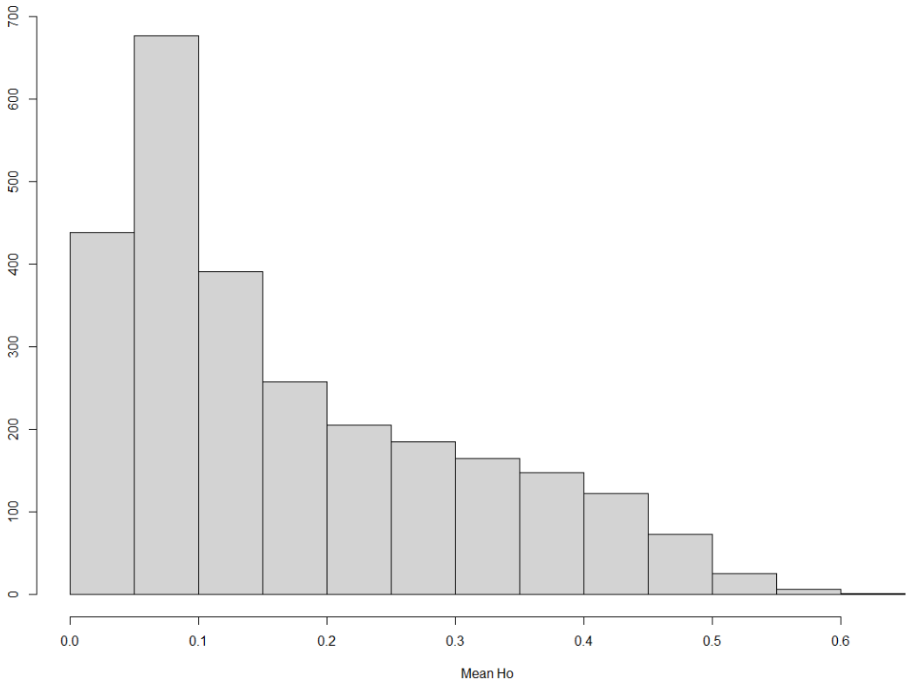


**Figure 1.** Mean observed heterozygosity (Ho) across the 2963 filtered SNP loci. Only 561 SNP loci (20.8%) had an average Ho greater than 0.3.

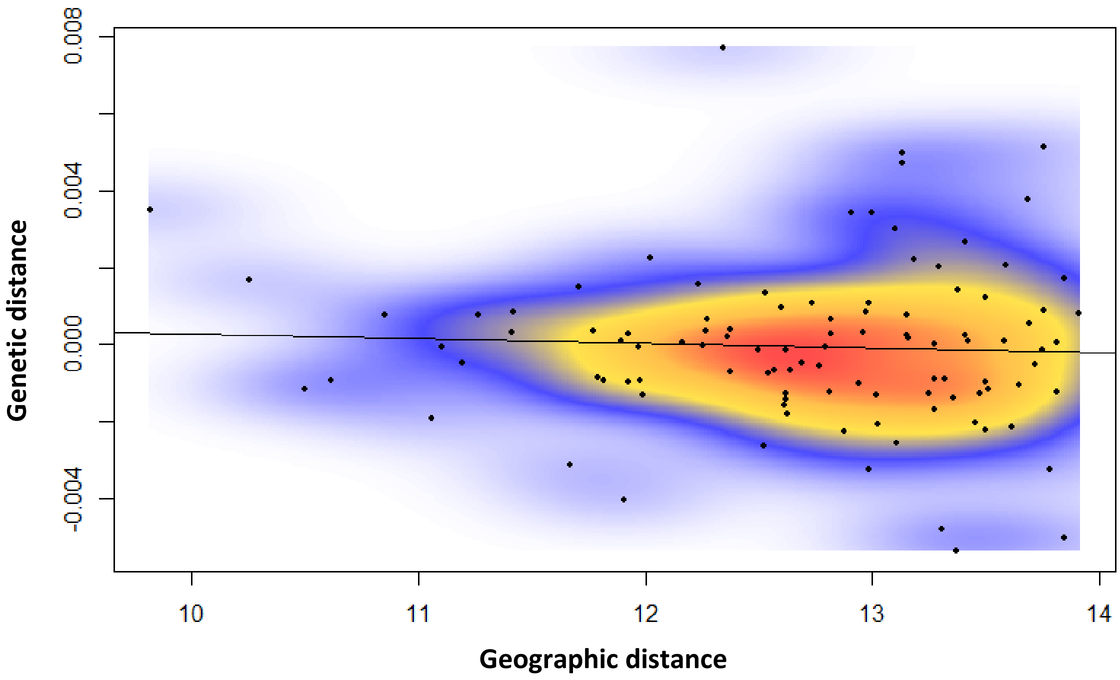


**Figure 2.** Isolation by distance with the 'ibd' function of the R package 'dartR' with 1000 permutations (r: -0.053, p = 0.685).

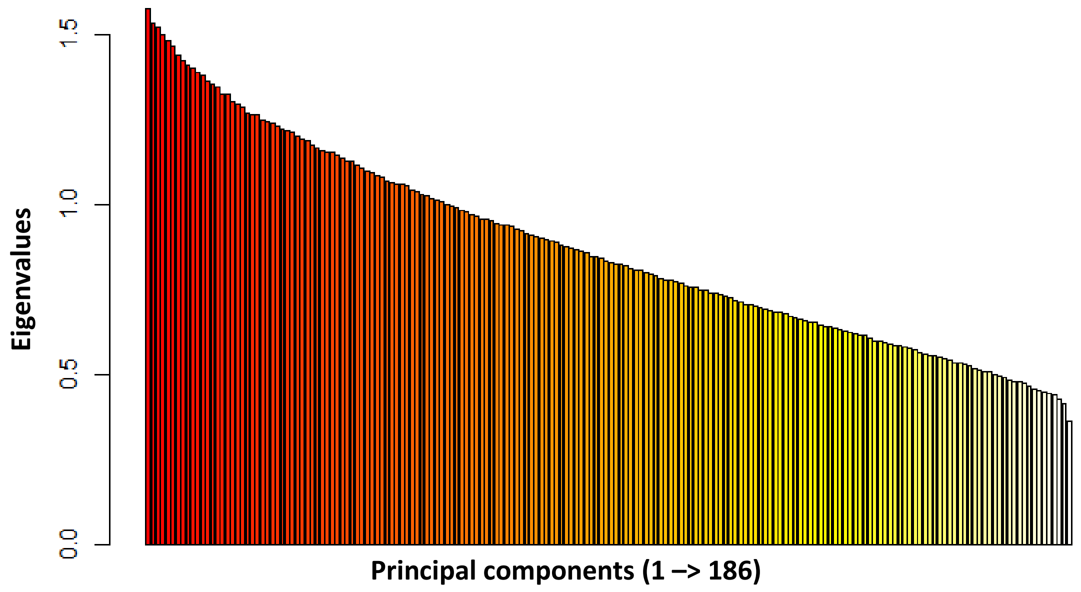


**Figure 3.** Eigenvalues which represent the amount of variation explained by each principal component, attained from the 'glPca' function of the R package 'adegenet'.

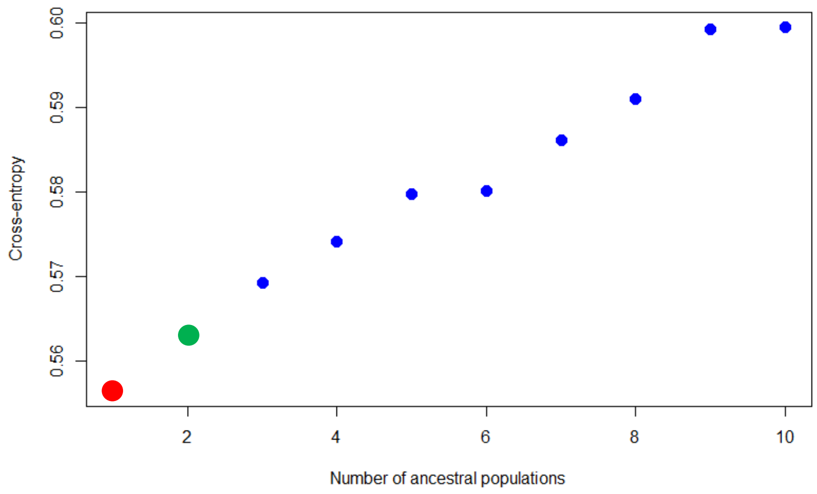


**Figure 4.** Minimal cross entropy for each number of ancestral populations (K) from 1 to 10, with 100 replications for each value of K. The value of K that best represents the population history is shown in red, and the value of K we used for the admixture proportions is shown in green. We chose to perform the analysis with a K value of 2 so that admixture proportions could be represented on a geographic map.

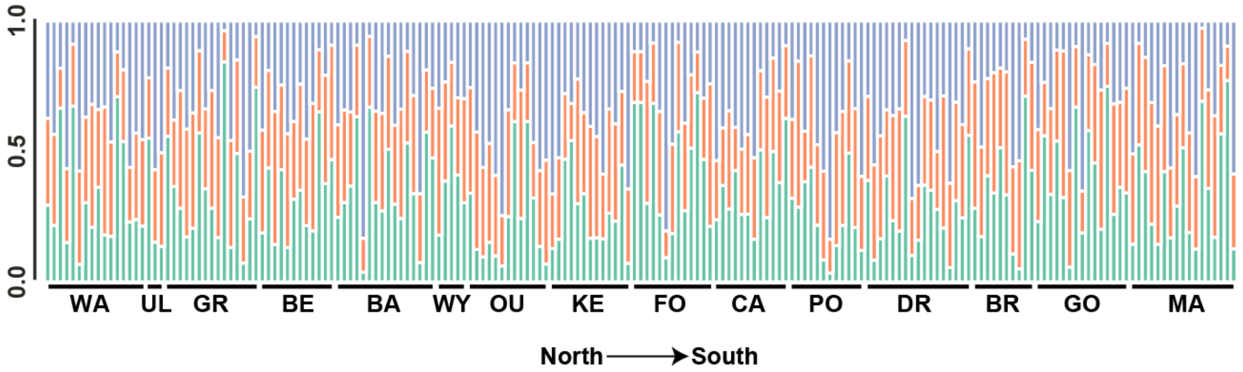


**Figure 5.** STRUCTURE analysis using IPYRAD filtered data. Populations were sorted geographically from north to south. To reinforce the analyses of the filtered SNP dataset, we also attained the raw FastQ files from DArTseq, and converted these into a VCF file with a custom ipyrad pipeline. We then filtered these VCF formatted data by removing any individuals with more than 50% missing data, any sites with more than 20% missing genotypes, and any sites with a minor allele frequency of less than 5%. This resulted in a dataset of 7,862 SNPs. We then analysed this ipyrad-filtered dataset using Structure v.2.3.4 (https://web.stanford.edu/group/pritchardlab/structure.html) across a K value range of 1 to 10 with 10 replications per K. An initial analysis was completed to determine the number of Markov Chain Monte Carlo (MCMC) iterations required to achieve stationarity in Fst and log(α) summary statistics for each K value. The optimal configuration was determined to be 100,000 iterations with 5,000 discarded as burn in and these parameters were used for all subsequent analyses. Following Structure analysis, the best K value for each data set was selected using the Evanno method implemented within the Structure Harvester online interface (available at: http://taylor0.biology.ucla.edu/structureHarvester/). To create plots comparable between analyses, results for each K value were permuted across all replicates using CLUMPP v1.1.2 and then plotted using Toyplot 0.18.0 (https://github.com/sandialabs/toyplot).

**Table 1.** Raw specimen data and locality information

| Specimen ID | Population | Latitude | Longitude | Sex |
|---|---|---|---|---|
| L0001-MtKeira_Female | MtKeira | -34.4047 | 150.8713 | Female |
| L0002-MtKeira_Female | MtKeira | -34.4047 | 150.8713 | Female |
| L0003-MtKeira_Female | MtKeira | -34.4047 | 150.8713 | Female |
| L0004-MtKeira_Female | MtKeira | -34.4047 | 150.8713 | Female |
| L0005-MtKeira_Female | MtKeira | -34.4047 | 150.8713 | Female |
| L0006-MtKeira_Female | MtKeira | -34.4047 | 150.8713 | Female |
| L0012-MtKeira_Male | MtKeira | -34.4047 | 150.8713 | Male |
| L0013-MtKeira_Male | MtKeira | -34.4047 | 150.8713 | Male |
| L0014-MtKeira_Male | MtKeira | -34.4047 | 150.8713 | Male |
| L0015-MtKeira_Male | MtKeira | -34.4047 | 150.8713 | Male |
| L0016-MtKeira_Male | MtKeira | -34.4047 | 150.8713 | Male |
| L0017-MtKeira_Male | MtKeira | -34.4047 | 150.8713 | Male |
| L0031-Foxground_Female | Foxground | -34.6982 | 150.792 | Female |
| L0032-Foxground_Female | Foxground | -34.6982 | 150.792 | Female |
| L0033-Foxground_Female | Foxground | -34.6982 | 150.792 | Female |
| L0034-Foxground_Female | Foxground | -34.6982 | 150.792 | Female |

| | | | | |
|---|---|---|---|---|
| L0035-Foxground_Female | Foxground | -34.6982 | 150.792 | Female |
| L0036-Foxground_Female | Foxground | -34.6982 | 150.792 | Female |
| L0041-Foxground_Male | Foxground | -34.6982 | 150.792 | Male |
| L0042-Foxground_Male | Foxground | -34.6982 | 150.792 | Male |
| L0043-Foxground_Male | Foxground | -34.6982 | 150.792 | Male |
| L0044-Foxground_Male | Foxground | -34.6982 | 150.792 | Male |
| L0045-Foxground_Male | Foxground | -34.6982 | 150.792 | Male |
| L0046-Foxground_Male | Foxground | -34.6982 | 150.792 | Male |
| L0074-MtCambewarra_Male | MtCambewarra | -34.8049 | 150.5722 | Male |
| L0075-MtCambewarra_Male | MtCambewarra | -34.8049 | 150.5722 | Male |
| L0076-MtCambewarra_Male | MtCambewarra | -34.8049 | 150.5722 | Male |
| L0077-MtCambewarra_Male | MtCambewarra | -34.8049 | 150.5722 | Male |
| L0078-MtCambewarra_Male | MtCambewarra | -34.8049 | 150.5722 | Male |
| L0079-MtCambewarra_Male | MtCambewarra | -34.8049 | 150.5722 | Male |
| L0097-MtCambewarra_Female | MtCambewarra | -34.8049 | 150.5722 | Female |
| L0098-MtCambewarra_Female | MtCambewarra | -34.8049 | 150.5722 | Female |
| L0099-MtCambewarra_Female | MtCambewarra | -34.8049 | 150.5722 | Female |
| L0100-MtCambewarra_Female | MtCambewarra | -34.8049 | 150.5722 | Female |
| L0101-MtCambewarra_Female | MtCambewarra | -34.8049 | 150.5722 | Female |
| L0102-MtCambewarra_Female | MtCambewarra | -34.8049 | 150.5722 | Female |
| L0115-PointerGap_Male | PointerGap | -35.2605 | 150.3582 | Male |
| L0116-PointerGap_Male | PointerGap | -35.2605 | 150.3582 | Male |
| L0117-PointerGap_Male | PointerGap | -35.2605 | 150.3582 | Male |
| L0118-PointerGap_Male | PointerGap | -35.2605 | 150.3582 | Male |
| L0119-PointerGap_Male | PointerGap | -35.2605 | 150.3582 | Male |
| L0120-PointerGap_Male | PointerGap | -35.2605 | 150.3582 | Male |
| L0142-PointerGap_Female | PointerGap | -35.2605 | 150.3582 | Female |
| L0143-PointerGap_Female | PointerGap | -35.2605 | 150.3582 | Female |
| L0144-PointerGap_Female | PointerGap | -35.2605 | 150.3582 | Female |
| L0145-PointerGap_Female | PointerGap | -35.2605 | 150.3582 | Female |
| L0146-PointerGap_Female | PointerGap | -35.2605 | 150.3582 | Female |
| L0147-PointerGap_Female | PointerGap | -35.2605 | 150.3582 | Female |
| L0151-BarringtonTops_Male | BarringtonTops | -32.1508 | 151.5248 | Male |
| L0152-BarringtonTops_Male | BarringtonTops | -32.1508 | 151.5248 | Male |
| L0153-BarringtonTops_Male | BarringtonTops | -32.1508 | 151.5248 | Male |
| L0154-BarringtonTops_Male | BarringtonTops | -32.1508 | 151.5248 | Male |
| L0155-BarringtonTops_Male | BarringtonTops | -32.1508 | 151.5248 | Male |
| L0156-BarringtonTops_Male | BarringtonTops | -32.1508 | 151.5248 | Male |
| L0157-BarringtonTops_Male | BarringtonTops | -32.1508 | 151.5248 | Male |
| L0158-BarringtonTops_Male | BarringtonTops | -32.1508 | 151.5248 | Male |
| L0171-BarringtonTops_Female | BarringtonTops | -32.1508 | 151.5248 | Female |
| L0172-BarringtonTops_Female | BarringtonTops | -32.1508 | 151.5248 | Female |
| L0173-BarringtonTops_Female | BarringtonTops | -32.1508 | 151.5248 | Female |
| L0174-BarringtonTops_Female | BarringtonTops | -32.1508 | 151.5248 | Female |

| | | | | |
|---|---|---|---|---|
| L0175-BarringtonTops_Female | BarringtonTops | -32.1508 | 151.5248 | Female |
| L0176-BarringtonTops_Female | BarringtonTops | -32.1508 | 151.5248 | Female |
| L0177-BarringtonTops_Female | BarringtonTops | -32.1508 | 151.5248 | Female |
| L0178-BarringtonTops_Female | BarringtonTops | -32.1508 | 151.5248 | Female |
| L0207-UlidarraNP_Male | UlidarraNP | -30.2488 | 153.0855 | Male |
| L0208-UlidarraNP_Male | UlidarraNP | -30.2488 | 153.0855 | Male |
| L0209-UlidarraNP_Male | UlidarraNP | -30.2488 | 153.0855 | Male |
| L0210-Bellangry_Male | Bellangry | -31.2892 | 152.537 | Male |
| L0211-Bellangry_Male | Bellangry | -31.2892 | 152.537 | Male |
| L0212-Bellangry_Male | Bellangry | -31.2892 | 152.537 | Male |
| L0213-Bellangry_Male | Bellangry | -31.2892 | 152.537 | Male |
| L0214-Bellangry_Male | Bellangry | -31.2892 | 152.537 | Male |
| L0215-Bellangry_Male | Bellangry | -31.2892 | 152.537 | Male |
| L0239-Bellangry_Male | Bellangry | -31.2892 | 152.537 | Male |
| L0240-Bellangry_Female | Bellangry | -31.2892 | 152.537 | Female |
| L0241-Bellangry_Female | Bellangry | -31.2892 | 152.537 | Female |
| L0242-Bellangry_Female | Bellangry | -31.2892 | 152.537 | Female |
| L0243-Bellangry_Female | Bellangry | -31.2892 | 152.537 | Female |
| L0244-Bellangry_Female | Bellangry | -31.2892 | 152.537 | Female |
| L0249-Ourimbah_Male | Ourimbah | -33.3649 | 151.3939 | Male |
| L0250-Ourimbah_Male | Ourimbah | -33.3649 | 151.3939 | Male |
| L0251-Ourimbah_Male | Ourimbah | -33.3649 | 151.3939 | Male |
| L0252-Ourimbah_Male | Ourimbah | -33.3649 | 151.3939 | Male |
| L0253-Ourimbah_Male | Ourimbah | -33.3649 | 151.3939 | Male |
| L0254-Ourimbah_Male | Ourimbah | -33.3649 | 151.3939 | Male |
| L0267-Ourimbah_Female | Ourimbah | -33.3649 | 151.3939 | Female |
| L0268-Ourimbah_Female | Ourimbah | -33.3649 | 151.3939 | Female |
| L0269-Ourimbah_Female | Ourimbah | -33.3649 | 151.3939 | Female |
| L0270-Ourimbah_Female | Ourimbah | -33.3649 | 151.3939 | Female |
| L0271-Ourimbah_Female | Ourimbah | -33.3649 | 151.3939 | Female |
| L0272-Ourimbah_Female | Ourimbah | -33.3649 | 151.3939 | Female |
| L0276-WyrrabalongNP_Male | WyrrabalongNP | -33.2938 | 151.5356 | Male |
| L0277-WyrrabalongNP_Male | WyrrabalongNP | -33.2938 | 151.5356 | Male |
| L0278-WyrrabalongNP_Male | WyrrabalongNP | -33.2938 | 151.5356 | Male |
| L0279-WyrrabalongNP_Male | WyrrabalongNP | -33.2938 | 151.5356 | Male |
| L0280-WyrrabalongNP_Male | WyrrabalongNP | -33.2938 | 151.5356 | Male |
| L0281-GrahamsTrail_Male | GrahamsTrail | -30.4245 | 152.8304 | Male |
| L0282-GrahamsTrail_Male | GrahamsTrail | -30.4245 | 152.8304 | Male |
| L0283-GrahamsTrail_Male | GrahamsTrail | -30.4245 | 152.8304 | Male |
| L0284-GrahamsTrail_Male | GrahamsTrail | -30.4245 | 152.8304 | Male |
| L0285-GrahamsTrail_Male | GrahamsTrail | -30.4245 | 152.8304 | Male |
| L0286-GrahamsTrail_Male | GrahamsTrail | -30.4245 | 152.8304 | Male |
| L0287-GrahamsTrail_Male | GrahamsTrail | -30.4245 | 152.8304 | Male |
| L0291-GrahamsTrail_Female | GrahamsTrail | -30.4245 | 152.8304 | Female |

| | | | | |
|---|---|---|---|---|
| L0292-GrahamsTrail_Female | GrahamsTrail | -30.4245 | 152.8304 | Female |
| L0293-GrahamsTrail_Female | GrahamsTrail | -30.4245 | 152.8304 | Female |
| L0294-GrahamsTrail_Female | GrahamsTrail | -30.4245 | 152.8304 | Female |
| L0295-GrahamsTrail_Female | GrahamsTrail | -30.4245 | 152.8304 | Female |
| L0296-GrahamsTrail_Female | GrahamsTrail | -30.4245 | 152.8304 | Female |
| L0297-GrahamsTrail_Female | GrahamsTrail | -30.4245 | 152.8304 | Female |
| L0298-Washpool_Male | Washpool | -29.47 | 152.316 | Male |
| L0299-Washpool_Male | Washpool | -29.47 | 152.316 | Male |
| L0300-Washpool_Male | Washpool | -29.47 | 152.316 | Male |
| L0301-Washpool_Male | Washpool | -29.47 | 152.316 | Male |
| L0302-Washpool_Male | Washpool | -29.47 | 152.316 | Male |
| L0304-Washpool_Male | Washpool | -29.47 | 152.316 | Male |
| L0305-Washpool_Male | Washpool | -29.47 | 152.316 | Male |
| L0306-Washpool_Male | Washpool | -29.47 | 152.316 | Male |
| L0327-Washpool_Female | Washpool | -29.47 | 152.316 | Female |
| L0328-Washpool_Female | Washpool | -29.47 | 152.316 | Female |
| L0329-Washpool_Female | Washpool | -29.47 | 152.316 | Female |
| L0330-Washpool_Female | Washpool | -29.47 | 152.316 | Female |
| L0331-Washpool_Female | Washpool | -29.47 | 152.316 | Female |
| L0332-Washpool_Female | Washpool | -29.47 | 152.316 | Female |
| L0333-Washpool_Female | Washpool | -29.47 | 152.316 | Female |
| L0334-Washpool_Female | Washpool | -29.47 | 152.316 | Female |
| L0335-MtDromedary_Male | MtDromedary | -36.2946 | 150.0337 | Male |
| L0336-MtDromedary_Male | MtDromedary | -36.2946 | 150.0337 | Male |
| L0337-MtDromedary_Male | MtDromedary | -36.2946 | 150.0337 | Male |
| L0338-MtDromedary_Male | MtDromedary | -36.2946 | 150.0337 | Male |
| L0339-MtDromedary_Male | MtDromedary | -36.2946 | 150.0337 | Male |
| L0340-MtDromedary_Male | MtDromedary | -36.2946 | 150.0337 | Male |
| L0341-MtDromedary_Male | MtDromedary | -36.2946 | 150.0337 | Male |
| L0342-MtDromedary_Male | MtDromedary | -36.2946 | 150.0337 | Male |
| L0350-MtDromedary_Female | MtDromedary | -36.2946 | 150.0337 | Female |
| L0351-MtDromedary_Female | MtDromedary | -36.2946 | 150.0337 | Female |
| L0352-MtDromedary_Female | MtDromedary | -36.2946 | 150.0337 | Female |
| L0353-MtDromedary_Female | MtDromedary | -36.2946 | 150.0337 | Female |
| L0354-MtDromedary_Female | MtDromedary | -36.2946 | 150.0337 | Female |
| L0355-MtDromedary_Female | MtDromedary | -36.2946 | 150.0337 | Female |
| L0356-MtDromedary_Female | MtDromedary | -36.2946 | 150.0337 | Female |
| L0357-MtDromedary_Female | MtDromedary | -36.2946 | 150.0337 | Female |
| L0358-MtDromedary_Female | MtDromedary | -36.2946 | 150.0337 | Female |
| L0361-BrownMountain_Male | BrownMountain | -36.5972 | 149.444 | Male |
| L0362-BrownMountain_Male | BrownMountain | -36.5972 | 149.444 | Male |
| L0363-BrownMountain_Male | BrownMountain | -36.5972 | 149.444 | Male |
| L0364-BrownMountain_Male | BrownMountain | -36.5972 | 149.444 | Male |
| L0365-BrownMountain_Male | BrownMountain | -36.5972 | 149.444 | Male |

| | | | | |
|---|---|---|---|---|
| L0366-BrownMountain_Male | BrownMountain | -36.5972 | 149.444 | Male |
| L0367-BrownMountain_Male | BrownMountain | -36.5972 | 149.444 | Male |
| L0368-BrownMountain_Male | BrownMountain | -36.5972 | 149.444 | Male |
| L0369-BrownMountain_Female | BrownMountain | -36.5972 | 149.444 | Male |
| L0370-BrownMountain_Female | BrownMountain | -36.5972 | 149.444 | Male |
| L0371-MaxwellsRainforest_Male | MaxwellsRainforest | -37.4145 | 149.8138 | Male |
| L0372-MaxwellsRainforest_Male | MaxwellsRainforest | -37.4145 | 149.8138 | Male |
| L0373-MaxwellsRainforest_Male | MaxwellsRainforest | -37.4145 | 149.8138 | Male |
| L0374-MaxwellsRainforest_Male | MaxwellsRainforest | -37.4145 | 149.8138 | Male |
| L0375-MaxwellsRainforest_Male | MaxwellsRainforest | -37.4145 | 149.8138 | Male |
| L0376-MaxwellsRainforest_Male | MaxwellsRainforest | -37.4145 | 149.8138 | Male |
| L0377-MaxwellsRainforest_Male | MaxwellsRainforest | -37.4145 | 149.8138 | Male |
| L0378-MaxwellsRainforest_Male | MaxwellsRainforest | -37.4145 | 149.8138 | Male |
| L0379-MaxwellsRainforest_Male | MaxwellsRainforest | -37.4145 | 149.8138 | Male |
| L0385-MaxwellsRainforest_Female | MaxwellsRainforest | -37.4145 | 149.8138 | Female |
| L0386-MaxwellsRainforest_Female | MaxwellsRainforest | -37.4145 | 149.8138 | Female |
| L0387-MaxwellsRainforest_Female | MaxwellsRainforest | -37.4145 | 149.8138 | Female |
| L0388-MaxwellsRainforest_Female | MaxwellsRainforest | -37.4145 | 149.8138 | Female |
| L0389-MaxwellsRainforest_Female | MaxwellsRainforest | -37.4145 | 149.8138 | Female |
| L0390-MaxwellsRainforest_Female | MaxwellsRainforest | -37.4145 | 149.8138 | Female |
| L0391-MaxwellsRainforest_Female | MaxwellsRainforest | -37.4145 | 149.8138 | Female |
| L0392-MaxwellsRainforest_Female | MaxwellsRainforest | -37.4145 | 149.8138 | Female |
| L0393-MaxwellsRainforest_Female | MaxwellsRainforest | -37.4145 | 149.8138 | Female |
| L0394-GoodeniaRainforest_Male | GoodeniaRainforest | -36.899 | 149.7154 | Male |
| L0395-GoodeniaRainforest_Male | GoodeniaRainforest | -36.899 | 149.7154 | Male |
| L0396-GoodeniaRainforest_Male | GoodeniaRainforest | -36.899 | 149.7154 | Male |
| L0397-GoodeniaRainforest_Male | GoodeniaRainforest | -36.899 | 149.7154 | Male |
| L0398-GoodeniaRainforest_Male | GoodeniaRainforest | -36.899 | 149.7154 | Male |
| L0399-GoodeniaRainforest_Male | GoodeniaRainforest | -36.899 | 149.7154 | Male |
| L0400-GoodeniaRainforest_Male | GoodeniaRainforest | -36.899 | 149.7154 | Male |
| L0401-GoodeniaRainforest_Male | GoodeniaRainforest | -36.899 | 149.7154 | Male |
| L0402-GoodeniaRainforest_Male | GoodeniaRainforest | -36.899 | 149.7154 | Male |
| L0411-GoodeniaRainforest_Female | GoodeniaRainforest | -36.899 | 149.7154 | Female |
| L0412-GoodeniaRainforest_Female | GoodeniaRainforest | -36.899 | 149.7154 | Female |
| L0413-GoodeniaRainforest_Female | GoodeniaRainforest | -36.899 | 149.7154 | Female |
| L0414-GoodeniaRainforest_Female | GoodeniaRainforest | -36.899 | 149.7154 | Female |
| L0415-GoodeniaRainforest_Female | GoodeniaRainforest | -36.899 | 149.7154 | Female |
| L0416-GoodeniaRainforest_Female | GoodeniaRainforest | -36.899 | 149.7154 | Female |
| L0417-GoodeniaRainforest_Female | GoodeniaRainforest | -36.899 | 149.7154 | Female |
